# 6mA-Sniper: Quantifying 6mA Sites in Eukaryotes at Single-Nucleotide Resolution

**DOI:** 10.1101/2023.03.16.530559

**Authors:** Jie Zhang, Qi Peng, Chengchuan Ma, Jiaxin Wang, Chunfu Xiao, Ting Li, Xiaoge Liu, Liankui Zhou, Wei-Zhen Zhou, Wanqiu Ding, Ni A. An, Li Zhang, Ying Liu, Chuan-Yun Li

## Abstract

While *N^6^*-methyldeoxyadenine (6mA) modification has been linked to fundamental regulatory processes in prokaryotes, its prevalence and functional implications in eukaryotes are controversial. Here, we report 6mA-Sniper to quantify 6mA sites in eukaryotes at single-nucleotide resolution. With 6mA-Sniper, we delineated an accurate 6mA profile in *C. elegans* with 2,034 sites, significantly enriched on sequences of [GC]G<u>A</u>G motif. Twenty-six of 39 6mA events with MnlI restriction endonuclease sites were experimentally verified, demonstrating the feasibility of this method. Notably, the enrichment of these 6mA sites on a specific sequence motif, their within-population conservation and the combinatorial patterns, and the selective constrains on them jointly support an active model for the shaping of the profile by some undiscovered methyltransferases. In a joint study (*Cell Research*, in revision), Ma *et al.* reported METL-9 as a new methyltransferase in shaping the basal and stress-dependent 6mA profile in *C. elegans*. Notably, with the 6mA profile identified by 6mA-Sniper at single-nucleotide resolution, we found that the levels of 6mAs are significantly decreased in strains with the removal of METL-9 (METL-9 KO-OP50), while generally increased after *P. aeruginosa* infection, further verified the efficiency of 6mA-Sniper in accurately pinpointing 6mA sites. Moreover, for the regions marked by 998 6mA sites emerged specifically after the infection, we identified an enrichment of the upregulated genes after the infection. The gene upregulations are likely mediated through a mutual exclusive crosstalk between 6mA and H3K27me3 modification, as supported by their co-occurrence, and the signal of increased H3K27me3 at regions marked by 6mAs depleted in METL-9 KO-OP50 strains. Notably, in different *C. elegans* strains, the cross-strain genetic variants removing 6mA sites are associated with the decreased expression of their host genes, and the removal of two randomly-selected 6mA events with genome editing directly decreased the expression of their host genes. We thus highlight 6mA regulation as a previously-neglected regulator of transcriptome in eukaryotes.

## INTRODUCTION

While 5-methylcytosine (5mC) modification has been linked to a series of fundamental regulatory mechanisms in eukaryotes (*1, 2*), the prevalence and functional significance of another type of DNA methylation in eukaryotes, 6mA (*N^6^*- methyladenine), remain to be addressed (*3, 4*). 6mA regulation was initially identified in prokaryotes with high abundance (*5*) and is linked to biological processes such as nucleosome positioning and transcription regulation (*6–8*). Several pilot studies with mass spectrometry or 6mA antibody-based approaches also reported the existence of 6mA regulation in eukaryotes (*9–11*). However, the reliability of these results was questioned, as contamination with high-level bacterial 6mA could confound the quantification by mass spectrometry or antibody-based immunoprecipitation (*12, 13*). Notably, a recent study found that the levels of 6mA in *C. elegans*, rodents, and primates are low or even undetectable after exclusion of the effects of bacterial contamination (*12*).

In contrast, sequence-based identification provides an efficient approach to control for bacterial contamination in 6mA quantification, through a specific focus on the targeted sequences of the species of interest. First, restriction enzyme-based 6mA identification could be used to verify candidate 6mA sites with specific recognition motifs, while it could not be used in unbiased delineation of genome-wide 6mA profiles due to the limited scope (*7, 14*). Second, DNA immunoprecipitation and sequencing (DIP-seq) with an anti-6mA antibody could be used to identify the 6mA profile at the genome-wide scale, but the findings are confounded by RNA m6A contamination due to nonspecific recognition by the antibodies (*4*). Third, third-generation sequencing methods, such as SMRT sequencing (*15*) and nanopore sequencing (*16*), provide an approach to directly detect 6mA signals. However, considering the high sequencing error rate and the large cross-molecule variations, an ultrahigh sequencing depth is needed to identify the 6mA sites with the standard analysis pipeline, which attempts to increase the signal-noise ratio by integrating the signals of different reads covering the same sites. The rate of false-positive identification is generally high, especially for eukaryotes with typically low 6mA abundance and thus insufficient cross-molecule signal integration. Notably, a recent study introduced SMRT circular consensus sequences with repeat measurements of interpulse duration (IPD) signals to control for cross-molecule variations, which successfully deconvolved global 6mA events from a genomic DNA sample and classified them by species of interest, genomic regions, and sources of contamination (*13*). However, the high cross-subreads variations in the IPD signals substantially limited its applications in pinpointing 6mA sites at single base resolution. Notably, this study did not report experimental verification of their candidate sites and the enrichment of these sites on some specific sequence motifs, and proposed deeper SMRT sequencing to achieve the precise mapping of specific 6mA events (*13*).

Notably, the identification of the dominant methyltransferase accounting for 6mA marks, or the “6mA writer”, could not only provide insight into the functional significance of this regulatory process, but provide a niche to evaluate the accuracy of the 6mA identification. Although several 6mA writers, such as Dam and EcoKI, have been identified in prokaryotes, the dominant methyltransferases accounting for 6mA profiles in eukaryotes are still to be addressed. As a note, DAMT-1 has been proposed as a 6mA methyltransferase in the regulation of mitochondrial stress and heat stress in *C. elegans* *(17-19)*, while further investigations with *in vitro* assays are needed to verify its 6mA methyltransferase activity, as well as its regulatory roles in shaping the base level of 6mA profiles in eukaryotes (*19*).

Overall, due to the difficulties in accurately identifying 6mA sites in eukaryotes, as well as the uncertainty of the dominant methyltransferases in shaping the ambiguous 6mA profile in eukaryotes, several studies have suggested that the 6mA modifications detected in eukaryote genomes may be the result of DNA polymerase misincorporation of recycled methylated nucleotides during DNA replication or repair, rather than direct DNA methylation by methyltransferases (*20–24*). In such a case, 6mA regulation, if present, may merely represent a misincorporation *via* the nucleotide-salvage pathway rather than an informative epigenetic mark with specific regulatory functions.

Here, we address these key issues through developing a third-generation sequencing- based method, 6mA-Sniper, to accurately quantify 6mA events in *C. elegans* at single- nucleotide resolution, which could be generalized to other eukaryotes such as yeast and mouse. Notably, in a joint study (*Cell Research*, in revision) (*25*), Ma *et al.* reported METL-9 as a new methyltransferase in shaping the basal and stress- dependent 6mA profile in *C. elegans*. With 6mA-Sniper, we further verified the dominant roles of METL-9 in shaping the 6mA profile at single-nucleotide resolution. Taken together, although we could not fully exclude the possibility of 6mA regulation as a process of DNA polymerase misincorporation of recycled methylated nucleotides, the enrichment of these 6mA sites on a specific sequence motif, their within-population conservation and the combinatorial patterns, the selective constrains on them, and the dominant contribution of a newly-identified methyltransferase METL-9 in shaping the base level and the stress-dependent changes of 6mA profiles jointly indicate 6mA regulation as an epigenetic regulator on eukaryote transcriptomes.

## RESULTS

### Identification of 6mA sites at single-nucleotide resolution

To identify 6mA sites from the third-generation sequencing signals, we first generated DNA samples of the positive control and negative control for 6mA sites. Briefly, five yeast DNA sequences covering most of the 256 types of 4-*mer* nucleotide combinations were amplified and subjected to an *in vitro* methyltransferase assay, with most of the A sites transformed to 6mAs. This sample was defined as a positive control for 6mA identification. A genome sample derived from yeast with genome-wide amplification was used as a negative control for these 6mA sites (**METHODS**). We then performed SMRT sequencing on the positive and negative control (Figure S1A, **METHODS**), respectively, and investigated the distributions of two kinetic parameters in SMRT sequencing, the value of the pulse fluorescence when synthesizing each nucleotide (the pulse width, PW) and the time between two adjacent incorporation events (the interpulse duration, IPD), of the two samples.

Notably, although the PW signals for the A sites were comparable between the positive and negative controls (Figure 1A), the IPD signals for the A sites were significantly higher in the positive control than in the negative control (Figure 1B, Wilcoxon test, P value < 2.2e-16). Consistent with this, a higher proportion of A sites was detected with high IPD scores (> 50) in the positive control sample, in contrast to the A sites in the negative control or other non-A sites in both the positive and negative control samples (Figure 1C). These findings thus indicate that the IPD signal, rather than the PW, is informative in differentiating normal A sites and 6mA sites in SMRT sequencing. Consistent with the previous findings, we detected large cross-molecule variations for the IPD score of each site as defined by different DNA molecules covering the same site (Figure S1B). Notably, when using SMRT circular consensus sequencing (CCS) for repeat measurement of the IPD signal of an individual DNA molecule, we also found high cross-subreads variations for the IPD scores of the same site, as defined by different rounds of sequencing for the same molecule (Figure 1D). Interestingly, in each round of SMRT sequencing, the changes in IPD scores for A sites showed a similar trend as those for the nearby sites, indicating that the varied microenvironment and the enzyme activity of the DNA polymerase in each round of SMRT sequencing of the same molecule contribute substantially to the cross-subreads variations for the IPD scores of the same site (Figure S1C). Overall, the large cross-molecule and cross-subreads variations thus hinder the identification of 6mA sites with direct comparisons of IPD signals of different sites.

**Figure 1.**
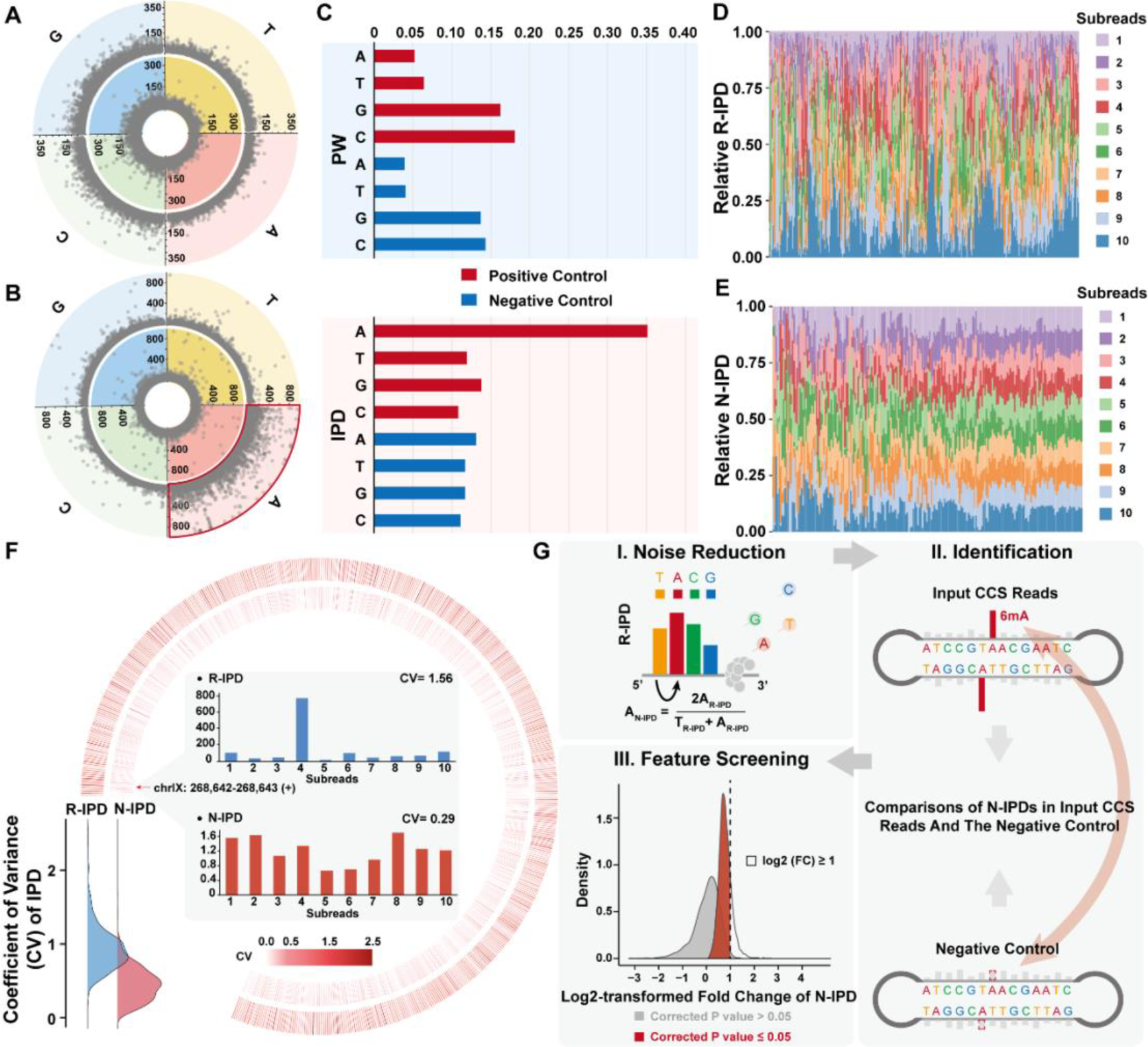
Identification of 6mA sites at single-nucleotide resolution. (**A**) Raw PW signals of the positive control (the second layer) and the negative control (the innermost layer). (**B**) Raw IPD signals of the positive control (the second layer) and negative control (the innermost layer). (**C**) The proportions of different sites with high IPD or PW signals (> 50) for the positive control and the negative control, respectively. (**D-E**) For each A site covered by ten subreads of one CCS reads (ID: 147,981,157; the plus strand), the raw IPD scores (**R-IPD**, **D**) and the noise reduction IPD scores (**N-IPD**, **E**) were normalized and shown. (**F**) The distribution and circular heatmaps showing the coefficient of variance of the R-IPD score (the second layer) and the N-IPD score (the innermost layer) for a CCS reads located on chrIX: 268,617- 270,510 (the plus strand). The R-IPD scores (upper panel) and N-IPD scores (lower panel) of one site (chrIX: 268,642-268,643, red arrow) on this CCS reads were also shown in the center of the circular heatmaps. (**G**) A workflow of the 6mA-Sniper in identifying 6mA sites at single-nucleotide resolution.

Promoted by the finding that the IPD score of the nearby sites is informative in deducing the status of SMRT sequencing at a specific temporospatial point and could possibly be used to decrease the signal-noise ratio in 6mA identification, we proposed a new method, 6mA-Sniper, to control for cross-molecule variations by introducing SMRT circular consensus sequencing for repeat measurement of the IPD signal of each individual DNA molecule, and especially to control for cross-subreads variations by normalizing the raw IPD score of each A site with that of the immediately upstream non-A site (N-IPD, noise reduction IPD, Figure 1E, **METHODS**). As expected, in comparison with the raw IPD signals typically used in previous protocols for calling 6mA sites, the N-IPD signals showed significantly decreased cross- subreads variations (Figure 1E and 1F, Wilcoxon test, P value < 2.2e-16).

We then identified 6mA sites at single nucleotide resolution on the basis of the N-IPD scores. Briefly, for each A site on the CCS reads, we calculated the N-IPD scores of this site based on the raw IPD scores of each round of SMRT sequencing covering this site and then compared the N-IPD scores with those calculated with all of the CCS reads in the negative control covering this site. The A sites showing significantly higher N-IPD scores than the negative control were selected as candidate 6mA sites (**METHODS**). As various *cis*- or *trans*-factors, such as other unknown DNA modifications and the unfavorable status of the DNA polymerase in sequencing low-complexity DNA sequences, may contribute to extreme changes in IPD signals and thus confound the comparisons, we further defined a threshold for the maximal N-IPD difference allowed between each candidate 6mA site and the negative control, according to the comparison of N-IPD distributions of A sites in the positive control and negative control. Briefly, an A site was defined as a 6mA site only when it showed significantly higher N-IPD scores than the negative control (Mann-Whitney test, Benjamini corrected P value ≤ 0.05), and the increased N-IPD score was located within a reasonable window (log2-transformed fold change < 1). The frequency of each 6mA site was then calculated on the basis of the portion of the CCS reads with 6mA modification on this site (Figure 1G).

### An accurate 6mA profile in *C. elegans*

As a proof of concept, we performed SMRT sequencing in two biological replicates of the *C. elegans* genome and one negative controls with a genome-wide amplified *C. elegans* genome (Table S1). A total of 1,089,796 and 319,864 CCS reads were generated for the *C. elegans* genome and the negative control, respectively, which were then subjected to 6mA-Sniper to identify 6mA sites. Overall, 2,034 sites supported by at least three positive CCS reads in both samples were identified as candidate 6mA sites in *C. elegans,* which were significantly enriched in sequence motif of [GC]G<u>A</u>G (Figure 2A and 2B). Consistently, the sequences with such a motif was also found in our joint study with Ma *et al.* as the most favorable substrates of the newly-identified 6mA methyltransferase in an *in vitro* transmethylation assay (*Cell Research, in revision*) (*25*). Notably, when identifying the candidate 6mA sites with the standard modification identification pipeline of SMRT sequencing, no specifically-enriched sequence motif was identified from the two *C. elegans* samples, in contrast to the negative control (Figure S2).

**Figure 2.**
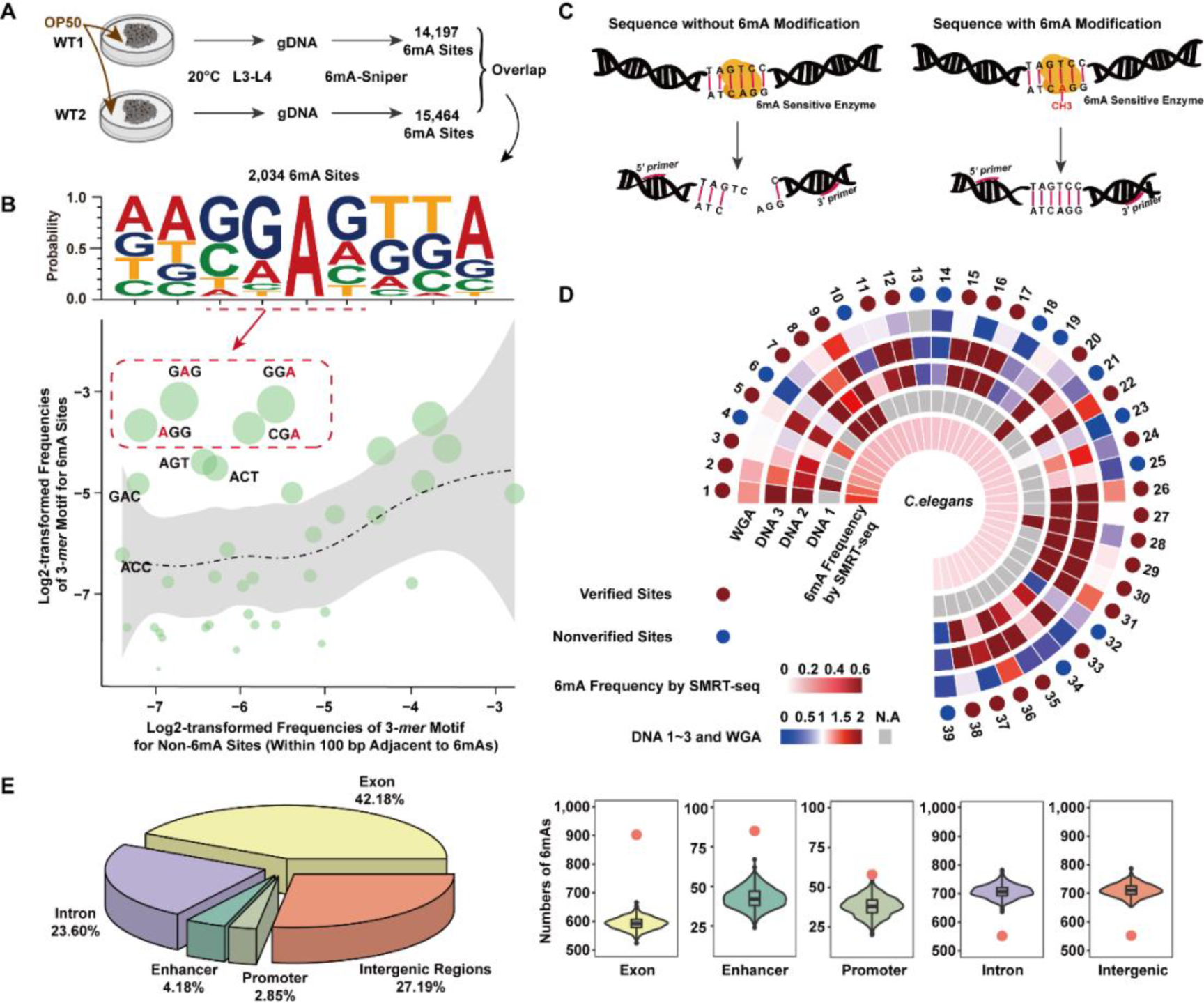
Identification of the 6mA profile in *C. elegans*. (**A**) A workflow for identifying the 6mA profile in wild-type *C. elegans* with 2,034 sites across the genome. (**B**) The sequence motifs of these 6mA sites are shown, with the most enriched 3-*mer* sequences centered by 6mA sites highlighted in red box. Each green bubble presents one of the various types of 3-*mer* nucleotide combinations with A sites. The size of each bubble represents the proportion of each 3-*mer* nucleotide combination, with the 6mA sites highlighted in red. The distribution of 3-*mer* nucleotide combination was estimated with random sequences, and shown in black dot line with gray regions representing 95% confidence intervals. (**C**) Schematic diagram showing the principle and procedures of the MSRE-qPCR experiment in the verification of candidate 6mA sites located on specific recognition motifs of restriction endonucleases. (**D**) A total of 39 6mA sites located on the GGAG motif were subjected to MnlI restriction enzyme assay. For each site, the frequencies of 6mA in three independent experiments, as well as in the negative control (**WGA**), were estimated and shown according to the color scale (**DNA 1**, **DNA 2**, **DNA 3**), with the missing data marked in gray. The frequency calculated by 6mA-Sniper was shown in the innermost layer according to the color scale. The verified and nonverified sites are marked in dark red and blue in the outermost layer, respectively. (**E**) The distribution of the 6mA sites on different genomic regions is shown. The distribution of random A sites on these genomic regions was estimated, with the real number of 6mA sites located on each genomic region highlighted in red points.

We then developed a restriction enzyme digestion assay to confirm that these sites represent *bona fide* 6mA events rather than technical artifacts. Among the full list of candidate 6mA events, we focused on a subset of them with the G<u>A</u>GG motif, which could be recognized by MnlI restriction endonuclease and thus be digested. Considering the sequences with G<u>6mA</u>GG could not be recognized by MnlI, the existence and frequency of the candidate 6mA sites could thus be determined through the estimation of the proportion of undigested DNA molecules (Figure 2C, **METHODS**). Among all of the 39 candidate 6mA sites feasible for testing with this assay (*e.g.*, located on nonrepetitive regions of the genome with the G<u>A</u>GG motif, and without other G<u>A</u>GG motif in the surrounding regions), 26 sites (or 66.7%) could be experimentally verified, showing a proportion of undigested molecules (or frequency of 6mA) significantly higher than that in the negative control (Figure 2D, **METHODS**). The sensitivity of this restriction enzyme digestion assay in quantifying 6mA events is relatively lower than that of the third-generation sequencing, leading to an underestimation of the verification rate. Notably, a substantial portion of the candidate sites not verified actually showed trends of higher proportion of undigested DNA molecules, while not distinguishable from the background at the strict threshold used in this study to control for false positives (Figure 2D). Overall, the enrichment of these candidate 6mA sites on specific sequence motif, as well as the experimental verifications with the restriction enzyme digestion assay, indicate that a large portion of these candidate sites should represent *bona fide* 6mA events in *C. elegans*.

To provide clues regarding the functional significance of this regulation, we further investigated the distribution of these 6mA events across the genome. Although these events are broadly distributed across the *C. elegans* genome, they are significantly enriched in regions with definite functions, such as the exonic (*Monte Carlo* test, P value < 0.001), promoter (P value < 0.001) and enhancer regions (P value < 0.001), while they are underrepresented in intronic (P value < 0.001) and intergenic regions (P value < 0.001, Figure 2E), indicating that the 6mA events may have some regulatory effects on the transcriptome.

### 6mA represents functional epigenetic mark shaped by methyltransferases

Recently, several studies have challenged the functional significance of 6mA regulation in eukaryotes and suggested that the weak signals of 6mA detected in eukaryotes represent a misincorporation *via* the nucleotide-salvage pathway (*20–24*). To test this hypothesis from the perspective of in-population conservation and the combination mode of the adjacent 6mA sites, we first identified a larger list of 6mA sites with 10,692 events in *C. elegans* using a relaxed threshold for the minimal number of positive CCS reads required (Figure S3A, **METHODS**). When comparing the 6mA sites identified by two independent samples, we found in-population conservation of the 6mA events. Briefly, a large portion of the 6mA sites identified in one sample could also be detected in another sample, especially for 6mA sites with higher frequencies (Figure 3A). The frequencies of the 6mA sites estimated using the positive CCS reads of the two samples were also positively correlated (Figure 3B). On the basis of the SMRT long reads, we further assessed the combinatorial emergence of adjacent 6mA sites on one molecule (Figure 3C). Notably, for the pairs of 6mA sites, the pattern of co-occurrence of the 6mA event on the same DNA molecule could be detected, especially for cases with the paired 6mA sites located within 100 bp (Figure 3D), a pattern previously reported in ADAR1 catalyzation (*26*).

**Figure 3.**
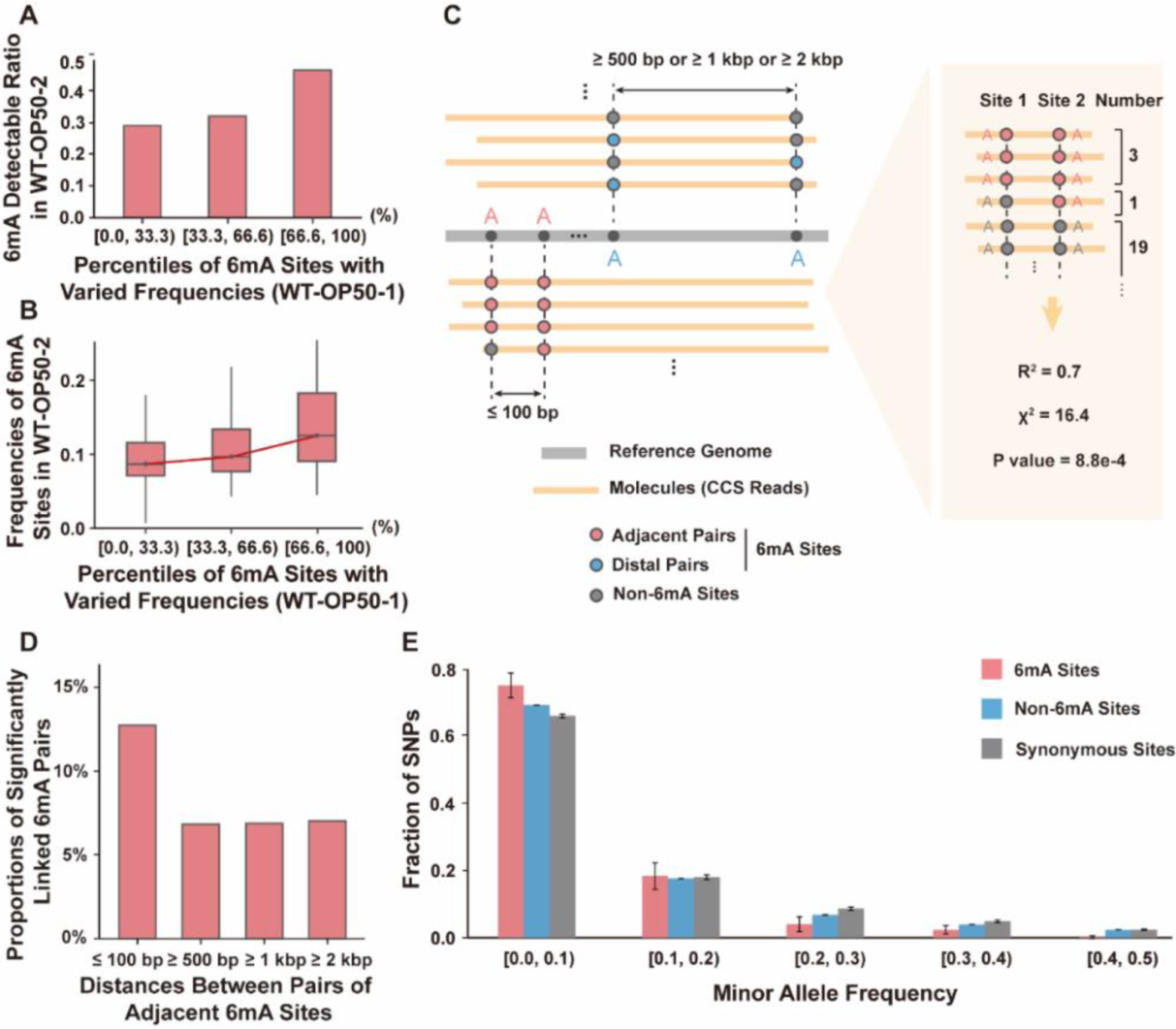
Conserved and selectively constrained 6mA profile in *C. elegans*. (**A, B**) For the 6mA sites identified in the first sample (**WT-OP50-1**), the portions of the 6mA sites detectable also in another sample (**WT-OP50-2**, **A**), and the distribution of the 6mA frequencies of these sites (**B**) were shown in three bins, according to the 6mA frequencies estimated by SMRT sequencing of the first sample. (**C**) Diagram showing the principle to quantify the combinatorial emergence of two adjacent 6mA sites (**Left panel**), and one case showing the steps of statistical analysis (**Right panel**). (**D**) For each pair of adjacent 6mA sites covered by at least 20 independent CCS reads, we tested whether they were significantly linked (**METHODS**). The pairs of 6mA sites were then divided into four bins according to their distances, and the proportions of significantly linked pairs were calculated and shown. (**E**) The distribution of minor allele frequency is shown for 6mA sites (red), non-6mA sites (blue) and synonymous sites as a neutral control (gray). The site frequency spectra of minor alleles are shown for different groups of sites, including the polymorphic sites occurring on 6mA events and not introducing nonsynonymous mutations (red), the polymorphic sites occurring on non-6mA sites and not introducing nonsynonymous mutations (blue), and the synonymous sites as a neutral control (gray). The site frequency spectrum for the minor allele was then estimated with 1,000 bootstraps to deduce the confidence intervals.

We further perform population genetics analysis to investigate whether these 6mA events represent functional regulations, or merely noise on DNA level during DNA replication or repair. In a population of 40 different *C. elegans* strains, we identified 77 polymorphic sites with the 6mA changed to other nucleotides across these strains, which lead to the removal of the 6mA modification in the corresponding strains (Figure S3B). When focusing on a portion of these polymorphic sites that removing the 6mAs but not introducing nonsynonymous mutations, we found that they were generally under selective constraints, in that the frequency spectra of the minor alleles showed an excess of low-frequency variants at these sites, in contrast to non-6mA sites (Wilcoxon one-tailed test, P value = 0.029), and the synonymous sites as a neutral control (Wilcoxon one-tailed test, P value = 0.012, Figure 3E). These findings thus indicate that the 6mA events are selectively constrained in general across the *C. elegans* strains, suggesting their functional significance in the worm.

Taken together, although we could not fully exclude the possibility of 6mA regulation as a misincorporation of recycled methylated nucleotides during DNA replication or repair, the within-population conservation of these sites, the co-occurrence of adjacent sites, the enriched sequence motif of these events, and the detection of selective constrains on these sites seem to support an active model for the shaping of the 6mA profile at specific chromosome locations by some undiscovered methyltransferases in *C. elegans*.

### Dominant contribution of METL-9 to the 6mA profiles in *C. elegans*

Promoted by the findings that the 6mA profile may be shaped by some undiscovered methyltransferases in *C. elegans*, we then attempted to identify the dominant “writer” of this new regulation. In a joint study with Ma *et al.* (*Cell Research, in revision*) (*25*), we identified METL-9 as the candidate methyltransferase accounting for the 6mA profiles in *C. elegans*, through a combinational strategy of functional screening and *in vitro* biochemical verifications for the methyltransferase activity. We then performed SMRT sequencing in a genetically modified *C. elegans* strain with METL-9 depletion (METL-9 KO-OP50), and delineated a 6mA profile with 1,384 sites with the combination of the SMRT sequencing data in both the wild-type (WT-OP50) and the METL-9 KO-OP50 samples (Figure 4A).

**Figure 4.**
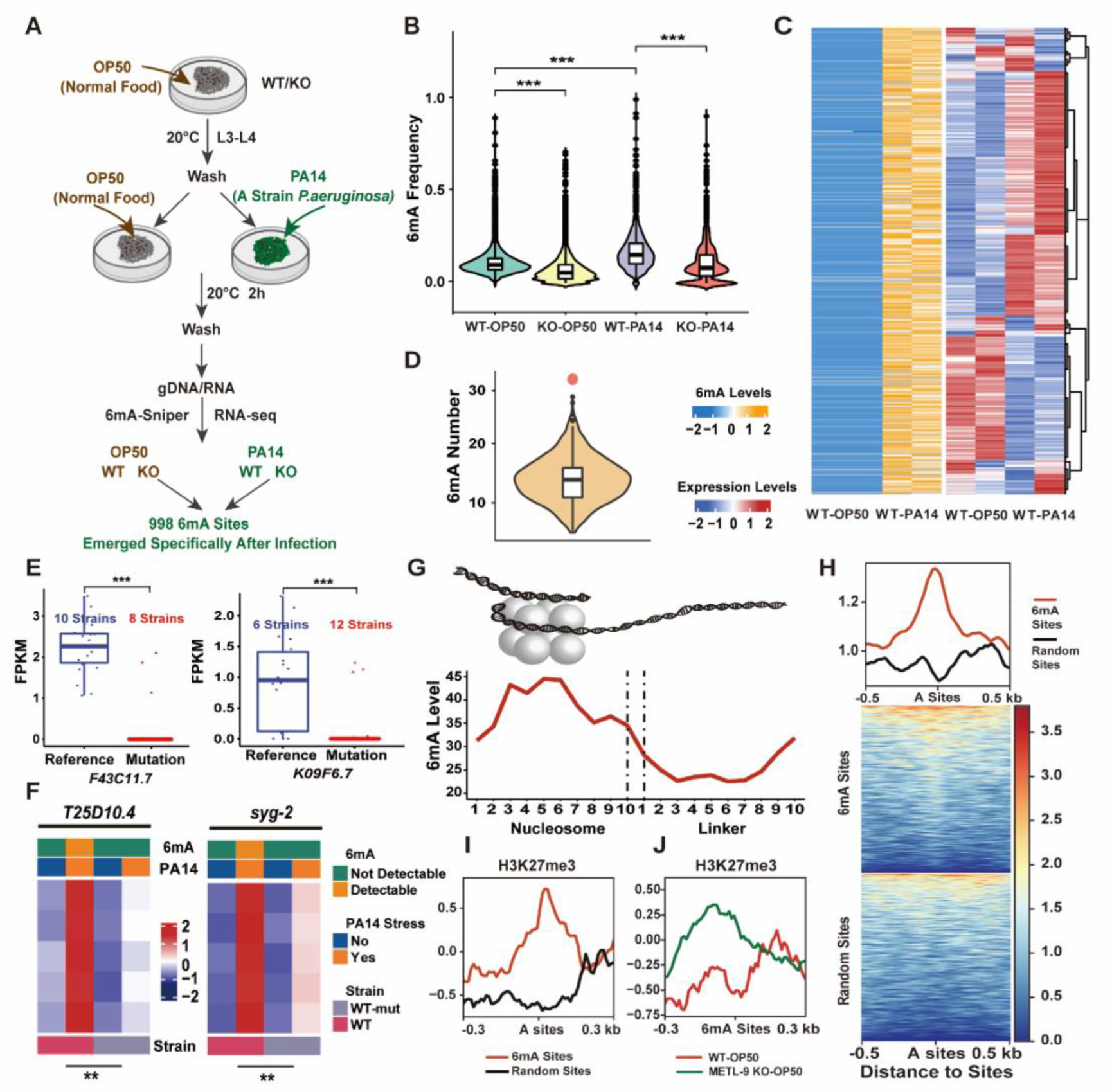
Regulation of 6mA events on the transcriptome. (**A**) A workflow for the investigation of the functions of METL-9 in *C. elegans*. (**B**) The distributions of the 6mA levels are shown for wild-type *C. elegans* (**WT-OP50**), *C. elegans* strains with METL-9 knockout (**METL-9 KO-OP50**), wild-type *C. elegans* strains treated with *P. aeruginosa* (**WT-PA14**), and **METL-9 KO-OP50** *C. elegans* strains treated with *P. aeruginosa* (**KO-PA14**). Wilcoxon test, ***P < 0.001. (**C**) For the newly-emerged 6mA sites after *P. aeruginosa* infection (left panel), the levels of 6mA are shown for two biological replicates of wild-type *C. elegans* (**WT-OP50**) and two biological replicates of the strains treated with *P. aeruginosa* (**WT-PA14**). For genes with at least one of these newly emerged 6mA sites, the expression levels of the genes in two biological replicates of **WT-OP50** and **WT-PA14** were normalized and shown in a heatmap according to the color scale. Only genes with FPKM > 2 in at least two of these samples were shown in the heatmap. (**D**) The number of 6mA sites located on upregulated genes after *P. aeruginosa* infection was shown in red point, together with the distribution of the background, as estimated using the same number of genes not differentially-expressed after *P. aeruginosa* infection. (**E**) The distributions of the gene expression in different *C. elegans* strains. For two polymorphic site co-localized with 6mA sites in the genic regions of the host gene, the expressions of the host gene were shown, for *C. elegans* strains with A allele at the polymorphic site (**Reference**), and the strains with other non-A alleles (**Mutation**). One-sided Wilcoxon tests were performed to test whether the expression levels between the two groups are significantly different. (**F**) Heatmaps showing the expression levels of *T25D10.4* and *syg-2*, in conditions with (**WT-PA14; WT-mut-PA14**) or without the *P. aeruginosa* treatment (**WT-OP50; WT-mut-OP50**). Two-tailed t-test. **P < 0.01. (**G**) The distribution of the 6mA sites across the nucleosome-located regions and the linker regions. (**H**) For the focal 6mA sites and their surrounding regions, the degrees of the nucleosome occupancy for these regions were aligned accordingly and shown in heatmap according to the color scale. (**I**) For the focal 6mA sites and their surrounding regions, the levels of H3K27me3 were shown (red line). The levels of H3K27me3 were also shown for randomly-selected A site and their surrounding regions as a background (black line). (**J**) For the 6mA events detected in **WT-OP50** but not in **METL-9 KO-OP50**, the levels of H3K27me3 at the focal 6mA site and its surrounding regions were shown, for *C. elegans* strains of **WT-OP50** (red line) and **METL 9 KO-OP50** (green line).

Notably, when comparing the 6mA profile between the wild-type and METL-9 KO- OP50 samples, we observed a significantly decreased 6mA level in the METL-9 KO- OP50 samples (Figure 4B, Wilcoxon test, P value < 2.2e-16). Moreover, in *C. elegans* treated with *P. aeruginosa*, we observed a globally-increased 6mA level at single-nucleotide resolution after the infection (Figure 4B, Wilcoxon test, P value < 2.2e-16), with the degree of change in well accordance with those as quantified by mass spectrometry in our joint study with Ma *et al.* (*Cell Research, in revision*) (*25*). In METL-9 KO-OP50, the 6mA levels were also increased after *P. aeruginosa* (PA14) infection, although the amplitude of the increasing was significantly lower (Figure 4B, Wilcoxon test, P value < 2.2e-16). Overall, these findings recapitulate the findings in Ma *et al.* that the methyltransferase METL-9 is one of the dominant writers in shaping the basal 6mA profiles in *C. elegans*, and largely accounts for the changed profiles after the *P. aeruginosa* infection, while other undiscovered methyltransferases should also have contributed to the profile.

### Regulatory roles of 6mA events on the transcriptome

Promoted by the findings of a joint study with Ma *et al.* (*Cell Research, in revision*) implicating 6mA into the transcriptional activation of genes associated with stress response, we then attempted to investigate whether 6mA could have some general regulatory effects on the transcriptome. Interestingly, for the list of 998 6mA sites newly-emerged in *C. elegans* treated with *P. aeruginosa* (Figure 4A), a large proportion of them are located on genes upregulated after the infection (Figure 4C). Accordingly, as for the whole set of genes significantly upregulated after *P. aeruginosa* infection, a significantly higher number of 6mA sites were located on them than the genomic background (Figure 4D, *Monte Carlo* test, P value < 0.001). These findings thus indicate the association between the 6mA events and the upregulation of the corresponding genes during the *P. aeruginosa* infection. To further clarify the causal relationship between them, we first investigated whether the cross-strain genetic variants removing the 6mA modification are associated with the decreased expression of the corresponding genes. Interestingly, in a population of 18 different *C. elegans* strains with the genome sequences and the matched transcriptome data, we identified 19 polymorphism sites occurring on the 6mA sites in genic regions, and ten of these sites are significantly associated with the differential expression of the corresponding genes. Interestingly, in 9 of these 10 sites (or 90%), the genes in strains with the genetic variants removing the 6mA sites are significantly downregulated (Figure 4E **and** Figure S4). These findings of eQTLs suggest that the 6mA events may positively regulate the expression of the corresponding genes.

Moreover, among the 998 6mA sites newly-emerged in *C. elegans* treated with *P. aeruginosa* (Figure 4A), we randomly selected two sites for further verification of their regulatory functions with causal relationship. For each 6mA event, we generated mutant *C. elegans* strains (WT-mut-OP50) from the WT-OP50, with the 6mA site replaced by other non-A base (Table S2). We then calculated the expression levels of the corresponding genes in WT-OP50 and WT-mut-OP50 strains, in conditions with the *P. aeruginosa* treatment (WT-PA14; WT-mut-PA14) or without the treatment (WT- OP50; WT-mut-OP50). Strikingly, for both of the newly-emerged 6mA events, the 6mA modification could be detected in WT-PA14, but not in WT-OP50, and theoretically not in WT-mut-OP50 and WT-mut-PA14. Accordingly, the expression levels of the host genes are comparable among WT-OP50, WT-mut-OP50 and WT-mut-PA14 (Figure 4F), but significantly upregulated in WT-PA14, the only conditions with the focused 6mA events detectable (Figure 4F). These lines of evidence thus support the notion that the 6mA events could positively regulate the expression of the host genes.

To further clarify the mechanism between the 6mA events and the upregulated expression of the corresponding genes, we investigated whether these 6mA events may crosstalk with other epigenetic regulations involved in transcription regulation. We first found that these 6mA events are enriched on nucleosomes in contrast to linker regions (Figure 4G and 4H), a pattern consistent with the nucleosome occupancy profile for 5mC events in mammals (*27*), and is in line with previous reports for increased levels of nucleosome occupancy and turnover rates in functional genomic regions (*28, 29*). Promoted by the finding of the enriched localization of 6mA-marked regions on nucleosomes, we then investigated whether these 6mA events may colocalize with genomic regions marked by major histone modifications, such as H3K4me2, H3K4me3 and H3K27me3 (Figure 4I, Figure S5). Interestingly, we found a significant overrepresentation of 6mA events in the regions marked by H3K27me3, a well-known epigenetic mark associated with the transcriptional inhibition (**METHODS**, Figure 4I). Consistently, for the regions marked by 6mA sites in WT-OP50 but not in METL-9 KO-OP50, we found a general increase of the H3K27me3 levels at these regions in METL-9 KO-OP50 strain (Figure 4J). These findings thus suggest the direct regulatory roles of 6mA events on gene expression, possibly through a mutual exclusive mechanism with H3K27me3 modification.

## DISCUSSION

Although the SMRT sequencing provides a theoretically efficient approach to directly detect 6mA signals, the high sequencing error rate and the large cross-molecule variations limited its applications. Recently, a new method was developed to estimate 6mA profiles across multiple species, using the CCS mode of SMRT sequencing to control for the cross-molecule variations (*13*). However, the high cross-subreads variations of the IPD signals, as indicated in Figure 1E in this study, substantially limited its application in pinpointing 6mA sites at single-nucleotide resolution. Here, the 6mA-Sniper we developed adequately controlled for both cross-molecule and cross-subread variations, and could thus be used to accurately quantify 6mA events at single-nucleotide resolution in eukaryotes (Figure 1). In a proof of concept study to identify 6mA sites in *C. elegans*, the significant enrichment of the candidate sites on specific sequence motifs, and verifications with restriction enzyme assays jointly demonstrated the accuracy of the new method. Notably, as stringent filtering criteria were used to control for false positives, the sensitivity of this new method in detecting 6mA events was not high, especially for those with low frequencies (Figure 2A and 2B). Overall, the accurate profiles of 6mA events at single-nucleotide resolution, although far from complete, could provide a representative niche to decipher the global functions of 6mA regulation in eukaryotes (Figure 2-4).

To investigate the feasibility of 6mA-Sniper in detecting 6mA sites in other species, we generated the SMRT sequencing data in yeast and the peripheral blood mononuclear cells (PBMCs) of mice (Table S1), and applied 6mA-Sniper to identify candidate 6mA events in these species. Overall, 31 6mA events on chromosome 6 in yeast, and 142 events on mitochondria genomes (mtDNA) in PBMCs of mice were identified. These candidate 6mA sites were also enriched in sequence motif of [GC]GAG, suggesting that they may similarly representing *bona fide* 6mA events (Figure S6), supporting the feasibility of 6mA-Sniper in detecting 6mA sites in other species, and the conserved features of 6mA regulation across distantly-related species.

Whether 6mA represents a functional epigenetic mark in eukaryotes remains controversial. Here, on the basis of the list of *bona fide* 6mA events identified in *C. elegans*, we found the enrichment of these 6mA sites on a specific sequence motif [GC]G<u>A</u>G, their within-population conservation, and the combinatorial emergence of adjacent 6mA sites, typically within 100 bp, on the same DNA molecule, a pattern previously reported in ADAR1-directed *trans*-regulations (*26*). Moreover, in a joint study with Ma *et al*. (*Cell Research, in revision*), we designed genome-wide screening assay to identify “writers” of the 6mA profile, and identified a new methyltransferase, METL-9, in shaping the 6mA profile in *C. elegans*. Interestingly, the *in vitro* transmethylation assay by Ma *et al*. indicates that the most favorable substrates of this methyltransferase is GG<u>A</u>G, consistent with the enriched sequence motif for the 6mA events identified in this study ([GC]G<u>A</u>G). Strikingly, the levels of 6mAs we identified at single-nucleotide resolution are significantly decreased in strains with the removal of METL-9, while generally increased after *P. aeruginosa* infection, clearly verifying METL-9 as the dominant methyltransferase in shaping the base level and stress-dependent changes of 6mA profile in *C. elegans*, and also verified the efficiency of 6mA-Sniper in accurately pinpointing 6mA sites at single-nucleotide resolution. Taken together, although we could not fully exclude the possibility of 6mA regulation as a process of DNA polymerase misincorporation of recycled methylated nucleotides, these findings, combined with the fact that these 6mA events are selectively constrained in general, jointly support the notion that at least a portion of these 6mA events should represent functional epigenetic mark shaped by methyltransferases (*20–24*).

While 6mA regulation has been linked to fundamental biological processes in prokaryotes, such as nucleosome positioning, DNA replication and transcription regulation, its functions in eukaryotes are largely controversial due to technical issues in accurately pinpointing 6mA events. Taking advantage of the accurate detection of 6mA sites at single-nucleotide resolution, we found that the 6mA events are selectively constrained in general across the *C. elegans* strains, suggesting their general functional significance in the worm (Figure 3).

Promoted by findings of a joint study with Ma *et al.* (*Cell Research, in revision*) implicating 6mA into the transcriptional activation of genes associated with stress response, we further investigated whether 6mA could have some general regulatory effects on the transcriptome. We observed a globally-increased 6mA level in *C. elegans* treated with *P. aeruginosa*, with 998 stress-dependent 6mA sites newly emerged after the infection (Figure 4B). The 6mA sites are preferentially located on nucleosomes, and more precisely, the regulatory regions marked by H3K27me3 (Figure 2D). Interestingly, in METL-9 KO-OP50 strains, we detected a globally- decreased level of 6mA events, while the levels of H3K27me3 are significantly increased at regions marked by the depleted 6mA events, indicating a mutual exclusive crosstalk between 6mA and H3K27me3 modification. We then verified the direct regulations of 6mA events on the expression of their host genes, through eQTLs analyses and CRISPR/Cas9-based gene editing of two randomly-selected 6mA events. These studies thus highlight 6mA regulation as a previously-neglected regulator of the transcriptome in eukaryotes.

## METHODS

### *C. elegans* cultivation, *P. aeruginosa* treatments, and DNA extraction

Animals of Bristol N2 (wild-type) and METL-9 KO-OP50 strains were maintained on nematode growth medium (NGM) at 20℃ and fed on *Escherichia coli* strain OP50. Animals at stage L4 were transferred to slow-killing plates with OP50 or *P. aeruginosa* (PA14) and fed for 2 hours at 20℃ before sample collection.

Animals were collected and washed with M9 buffer (42.3 mM Na_2_HPO_4_, 22.1 mM KH_2_PO_4_, 85.5 mM NaCl). The compact worm slurry was lysed in DNA lysis buffer (200 mM NaCl, 100 mM Tris-HCl pH 8.5, 50 mM EDTA (pH 8.0), 0.5% SDS) with 0.1 mg/mL proteinase K (Thermo) at 55℃ until clear. The samples were incubated with 0.5 mg/mL RNase A (Tiangen) at 37°C for 1 hour. The DNA was then isolated by phenol:chloroform:isopentanol (25:24:1) extraction and isopropanol precipitation. The DNA pellet was washed with 70% ethanol, dissolved in TE buffer, and incubated at 37°C for 1 hour with RNase A/T1 (Thermo, 1/20 volume) and RNase H (NEB, 1/50 volume). The DNA was then purified again.

### Yeast cultivation and DNA extraction

*Saccharomyces cerevisiae* strain W303 was cultured in liquid YPD to OD600 ≥ 1.0, which was then centrifuged at 2,500 rpm for 5 min at 4°C. The pellet was resuspended in 1 ml pre-cooled ddH_2_O and centrifuged at 8,000 rpm for 10 s at 4°C. The supernatant was removed, and the pellet was then vortexed with the remaining solution to resuspend the cells. The samples were vortexed with 200 μL phenol:chloroform:isopentanol (25:24:1), 0.4 g glass beads and 200 μL lysis buffer for 5 min. Then 200 μL 1X TE buffer was added, followed by centrifuging at 14,000 rpm for 5 min at 4°C. The supernatant was transferred to a new tube, and 450 μL chloroform:isopentanol (24:1) was added, followed by centrifuging at 14,000 rpm for 5 min at 4°C. The supernatant was the mixed with 1 mL pre-cooled 100% EtOH gently, and then centrifuged at 14,000 rpm for 5 min at 4°C. The supernatant was then removed, and the pellet was resuspended with 400 μL 1X TE buffer, and digested with 2 μL 20 mg/mL RnaseA at 37°C for 1 hour. After the digestion, 1 mL pre-cooled 100% EtOH and 10 μL 4 M acetic acid were added, followed by centrifuging at 14,000 rpm for 5 min at 4°C. The precipitation was then washed using 1 mL pre-cooled 100% EtOH for twice and dissolved in 1X TE buffer.

### Isolation of peripheral blood mononuclear cells of mice and DNA extraction

Female C57BL/6J mice were purchased from the Institutional Animal Care of Peking University. All animal studies were approved by the Institutional Animal Care and Use Committee of Peking University. 8 weeks old animals were used in the experiments. Peripheral blood mononuclear cells (PBMCs) were isolated from peripheral blood of these mice, using density gradient medium Pancoll (Solarbio). The obtained PBMCs were then washed by cold PBS for three times. The cell pellet was lysed in DNA lysis buffer with 0.1 mg/mL proteinase K at 55 ℃ until clear. The total DNA was then extracted as mentioned above.

### Construction of the positive and negative controls, library preparation and SMRT sequencing

To generate a positive control in 6mA identification, five DNA fragments of 2,000 bp with the most diverse distribution of 4-*mer* sequences were selected from the yeast genome (strain: S288c), amplified and inserted into the plasmid vector, and then methylated by EcoGII methyltransferase *in vitro*. Briefly, 0.5 μg plasmid was incubated with 5 U EcoGII with 160 μM SAM at 37 °C for 2 hours. The plasmid was then purified by phenol:chloroform:isopentanol (25:24:1) extraction and isopropanol precipitation. To generate a negative control for 6mA identification, the purified genomic DNA of wild-type *C. elegans* was subjected to whole genome amplification using the QIAGEN REPLI-g Mini Kit by following the instructions.

To determine the overall levels of the 6mA modification in the positive control, 50 ng of the plasmid was digested by 1 μL DNase I in a 20 μL reaction at 37°C for 8 hours, denatured at 95°C for 5 min, chilled on ice and then incubated with 1/10 volume of 100 mM NH_4_OAc (pH 5.3) and 1 μL of nuclease P1 overnight at 42°C. Then, 1/10 volume of 1 M NH_4_HCO_3_ and 1 μL of rSAP were added, and the reaction was incubated at 37°C for 8 hrs. The final product was centrifuged at 15,000 rpm for 30 min. LC‒MS/MS measurements of 2′-deoxyadenosine and *N^6^*-methyl-2′- deoxyadenosine were then conducted on a Waters I-Class UPLC coupled with a Xevo TQ-S microMass Spectrometer equipped with an electrospray ionization source (Waters Technologies, Milford, USA). Briefly, 1 μL digested DNA sample was injected into a Waters ACQUITY UPLC HSS T3, 2.1 mm × 100 mm, 1.8 μm column (Waters Technologies, Milford, USA) for separation, with the column temperature controlled at 30 ℃. Mobile phase A was H_2_O containing 0.1% formic acid, and mobile phase B was acetonitrile and 0.1% formic acid. The flow rate was set at 0.4 mL/min. During the whole analysis, the gradient was 0-1 min, 0% B; 1.5 min, 3% B; 2 min, 5% B; 3-5 min, 8% B; 5-6.5 min, 100% B; and 6.5-8 min, 0% B. The autosampler temperature was maintained at 4 ℃ to avoid sample degradation. The MRM detection of target compounds was performed in positive electrospray ionization (ESI+) mode. For the transition of m/z 252.0→136.0 (2′-deoxyadenosine), the cone voltage was 16 V, and the collision energy was 10 V; for the transition of m/z 266.0→150.0 (N6-methyl-2′-deoxyadenosine), the values were 8 V and 10 V, respectively.

The DNA samples were quantified by Qubit (Invitrogen) and examined by an Agilent 4,200 for integrity. The samples were then sheared by g-tube for three times. DNA fragments with lengths in the range of 1,000 bp-8,000 bp were selected for SMRT sequencing, in which the SMRTbell libraries were constructed according to the standard PacBio protocols, and the well-prepared libraries were sequenced by PacBio Sequel II with the CCS mode.

### Identification of 6mA sites at single nucleotide resolution

We developed a new method, 6mA-Sniper, to identify 6mA sites at single-nucleotide resolution. First, the raw SMRT sequencing reads were mapped to the reference genome of the species of interest using pbmm2 (version 1.1.0, *-c* 80 and *-l* 1000). A reference-free alignment among subreads of each CCS reads was then performed, and only CCS reads with ≥ 5 passes of subreads were kept for the following analyses. For each A site, the raw IPD signals (R-IPD) were extracted by pysam (version 0.15.4), which was then normalized with the IPD signals of the immediately upstream non-adenine site (N-IPD, noise reduction IPD, Figure 1E) to increase the signal-noise ratio. The effects of the noise reduction were then evaluated through the calculation of the coefficient of variation on the basis of the R-IPD scores or N-IPD scores. For each A site on each CCS reads, we then compared the N-IPD scores derived from the subreads of the sample of each round of SMRT sequencing covering the A site with those calculated with all of the CCS reads in the negative control covering the A site. A Mann-Whitney test was performed to test the difference in the two groups of N-IPD scores, with a Bonferroni corrected P value threshold of 0.05. The A site showed significantly higher N-IPD scores than the negative control and was then defined as the candidate 6mA site. We further defined a threshold for the maximal N-IPD difference allowed between each candidate 6mA site and the negative control, according to the comparison of N-IPD distributions of A sites between the positive and the negative control. Only candidate 6mA sites with increased N-IPD scores within a reasonable window (log2-transformed fold change < 1) were included. The frequency of each 6mA site was then estimated on the basis of the portion of the CCS reads with 6mA modification on this site.

On the basis of 6mA-Sniper, we delineated an accurate 6mA profile in *C. elegans* (ce11). Briefly, we performed SMRT sequencing in two biological replicates (WT- OP50-1 and WT-OP50-2) of the *C. elegans* genome and one negative controls with a genome-wide amplified *C. elegans* genome. A total of 1,089,796 and 319,864 CCS reads were generated for the *C. elegans* genome and the negative controls, respectively, which were subjected to 6mA-Sniper to identify 6mA sites. Overall, 2,034 sites supported by at least three positive CCS reads in both samples were identified as candidate 6mA sites in *C. elegans*.

We also applied 6mA-Sniper to identify candidate 6mA events on chromosome 6 of yeast DNA and mice PBMC mtDNA. The data preprocessing were performed following the pipeline as described above with some minor modifications. Briefly, considering the low 6mA level in yeast and mice, we used Benjamini-Hochberg test, rather than the Bonferroni test in the original pipeline, to pinpoint A sites with significantly higher N-IPD scores than the negative control, with a corrected P value threshold of 0.05. Moreover, considering the variations of the sequencing depths between DNA and mtDNA, a minimal proportion of 20% positive CCS reads for DNA, or 2% for mtDNA, are required to define a candidate 6mA site, respectively. Finally, 31 6mA sites were identified on chromosome 6 of yeast (sacCer3) DNA, and 142 6mA sites on mtDNA of mice PBMCs (mm10).

### MSRE-qPCR verification of candidate 6mA sites

100 ng DNA was treated with 0.5 μL xonuclease VII (Thermo) at 37°C for 1 hour, followed by 65°C for 20 min to inactivate the enzyme. Then, 0.5 μL MnlI (NEB) was added to the sample, which was incubated at 37°C for 2 hours and 85°C for 10 min. Then, 5 ng input DNA was subjected to qRT-PCR assays to determine the frequency of the candidate 6mA sites through the estimation of the proportion of undigested DNA molecules.

### Features of the 6mA profile in *C. elegans*

The motif analyses were performed with WebLogo3 (*30*). An inhouse script was developed to identify the enriched 3-*mer* sequences centered by 6mA sites, in contrast to random sequences as the negative control. When investigating the distributions of 6mA sites across the genome, the exonic, intronic and intergenic regions were defined according to the annotations (WBcel235.87). The enhancer and promoter regions in *C. elegans* were defined according to the regulatory annotations of Jänes *et al.* (*31*). The cumulative signals were calculated by deepTools (version 3.5.1) (*32*).

### Within-population conservation and the combination mode of 6mA sites

Considering the low abundance of 6mA regulation, to study its within-population conservation, we first identified a larger list of 6mA sites in *C. elegans* using a relaxed threshold, in which a threshold of two positive CCS reads, rather than three in the original pipeline, was used to define a candidate 6mA site. We first separately identified the 6mA sites in the two independent samples used in this study (WT-OP50- 1 and WT-OP50-2). For each 6mA site identified in WT-OP50-1, we then checked whether the site was also supported by positive CCS read in WT-OP50-2 SMRT sequencing. The 6mA sites were then divided into three bins according to their 6mA frequencies as defined in WT-OP50-1, and the proportion of 6mA sites also detectable in WT-OP50-2 was calculated and compared. We then focused on the shared 6mA sites (10,692 sites) identified separately in the two samples, and compared the frequencies of the 6mA sites as estimated separately by the proportions of positive reads in the two samples.

On the basis of the long reads generated in SMRT sequencing, we also assessed the combined emergence of the adjacent 6mA sites on one molecule. Briefly, for each pair of adjacent 6mA sites covered by at least 20 independent CCS reads, the frequencies of the four different combinations, including 6mA-6mA (*f_AA_*), 6mA-A (*f_Aa_*), A-6Ma (*f_aA_*) and A-A (*f_aa_*), were calculated on the basis of the CCS reads. The *D* value and the correlation coefficient *R^2^* were then defined as follows. A chi-square test was then performed to test whether the two 6mA events were significantly linked (M refers to the depth of the reads coverage; 1 and 2 refer to two adjacent sites, respectively.), with a Benjamini-Hochberg corrected P value threshold of 0.05. The pairs of 6mA sites were then divided into four bins according to their distances, and the proportions of significantly linked pairs were calculated and are shown.

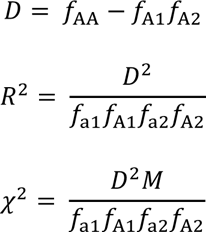

### Population genetics analyses

To investigate whether the 6mA profile is maintained by natural selection in *C. elegans*, we performed population genetic analyses in a population of 40 *C. elegans* strains (*33*). We identified 77 polymorphic sites with 6mA replaced with other nucleotides, leading to direct removal of the 6mA modification in the corresponding *C. elegans* strains. For each of these sites, the minor allele was defined with VCFtools (version 0.1.16). The site frequency spectra of minor alleles were then estimated for different groups of sites, including the polymorphic sites occurring on 6mA events but not introducing nonsynonymous mutations, the polymorphic sites occurring on non- 6mA sites but not introducing nonsynonymous mutations, and the synonymous sites as a neutral control. The site frequency spectrum for the minor allele was then estimated with 1,000 bootstraps to deduce the confidence intervals. Wilcoxon tests were then performed to compare the average minor allele frequency between different groups, with a threshold P value of 0.05.

### Effects of METL-9 KO-OP50 and *P. aeruginosa* infection in *C. elegans*

METL-9 KO-OP50 strains were provided by Dr. Ying Liu, and the information for the generation of this strain is described in detail in a joint study with Ma *et al*. (*Cell Research*, in revision) (*25*). The *P. aeruginosa* treatment was performed as described above. The METL-9 KO-OP50 samples and *P. aeruginosa* infection samples were prepared and subjected to the third-generation sequencing following the pipeline as mentioned above. Due to the relatively low sequencing depth for each sample, to control for the sampling bias in comparing the 6mA levels, we first combined the SMRT sequencing data of wild-type and METL-9 KO-OP50 samples to identify a reference list of 6mA sites with 16,365 sites, with each site supported by at least five positive CCS reads. Then, for each 6mA site, the frequency was calculated separately for each samples, with the sites of low coverage (reads coverage < 10) assigned as missing data in the corresponding sample.

To investigate the correlation between 6mA regulation and the expression of their host genes, we then quantified the transcriptome of these samples with poly(A)-enriched RNA-seq. The RNA extraction was performed by washing the worm pellets with M9 buffer and resuspending with TRIzol, then the samples were frozen in liquid nitrogen and homogenized. Chloroform extraction and isopropanol precipitation was used to isolate the total RNA. Two biological replicates of wild-type *C. elegans* and wild-type *C. elegans* treated with *P. aeruginosa* were then subject to deep sequencing on Illumina Platform with 150 bp paired-end mode. The RNA-seq reads were mapped to the *C. elegans* genome (ce11) using HISAT2 (version 2.2.1) (*34*) with default parameters, with low-quality reads removed by samtools (v1.12, -f 34 -F 1564 or -f 18-F 1580) (*35*) and sambamba (v0.8.2) (*36*). RNA-seq reads mapped on the genic regions were quantified and annotated using Subread feature Counts (version 2.0.2) (*37*). Gene expression levels were then calculated as FPKM values using edgeR (version 3.34.1) (*38*), and the differentially-expressed genes were identified using DESeq2 (version 1.32.0) (*39*), in which genes with expression fold change > 1.5 and an adjusted P value < 0.05 were defined as differentially expressed genes, and the genes with the difference of expression < 5% between wild-type and the sample with *P. aeruginosa* treatment were defined as genes not differentially-expressed. The number of 6mA sites located on upregulated genes after *P. aeruginosa* infection was then calculated, together with the distribution of the background, as estimated using the same number of genes not differentially-expressed for 1,000 times of bootstraps which was used to test whether the 6mA sites were significantly enriched on upregulated genes after *P. aeruginosa* infection. The public RNA-seq data of different *C. elegans* strains were also obtained from GSE186719 (*40*) for a cross-strain comparison of the expressions for 6mA-associated genes, using the identical pipeline as described above.

To verify the regulatory functions of 6mA events on the expression of their host genes with a causal relationship, we then developed two CRISPR/Cas9 genome editing strains of *C. elegans*, including the PHX6478 syg-2 (syb6478) strain with the 6mA site on *syg-2* gene (chrX:14660859 (-)) edited from A to T, and the PHX6484 T25D10.4 (syb6484) strain with the 6mA site on K10D11.5 edited (chrIV:12989886 (-)) edited from A to C. The animals with genome editing were outcrossed to the wild-type strain (N2), and the editings were verified by Sanger sequencing. Then, the wild-type and mutant strains of animals at stage L4 were transferred to slow-killing plates with OP50 or *P. aeruginosa* (PA14), and fed for 2 hours at 20℃ before sample collection. Total RNAs of these samples were then extracted using the pipeline as described above, which were further subjected to cDNA synthesis and RT-qPCR using SYBR GREEN PCR Master Mix, with the primers ACGATTATCAAACGCGGAGC (Forward) CTCGAACTCCCATGTTCGGA (Reverse) for *syg-2* and CGATTTGGACCGAGAAGGGT (Forward) CTGCTTGGCGAGTGAGTCTT (Reverse) for *T25D10.4*, respectively.

### MNase-seq analyses

L4 animals were collected and washed by M9 buffer. 500μL compacted worm slurry was homogenized in 2 mL Dounce Buffer (0.35 M sucrose, 15 mM HEPES pH 7.5, 0.5 mM EGTA, 0.5 mM MgCl2, 10 mM KCl, 0.1 mM EDTA, 1 mM dithiothreitol, 0.5% Triton X-100) with protease inhibitors. The mixture was spun at 150 g for 1 min at 4°C and the supernatant containing nuclei was transferred to a new tube (repeated for another round). Nuclei was then collected from supernatant by centrifugation at 4,000 g for 10min at 4°C, and resuspended in MNase reaction with 1mM CaCl2 and 25U/μL MNase (NEB, M0247S). The reaction was incubated at 16°C for 12min. Then equal volume of worm DNA lysis buffer with 2 mg/mL proteinase K was added and incubated at 60°C for 45 min with gentle shake. DNA was extracted and ethanol precipitated. 1/50 volume of RNase A/T1 was added and incubated at 37°C for 30min. DNA was extracted and ethanol precipitated, and the mono-nucleosome DNA fragments (150 – 200 bp) were separated by 2% agarose gel, and further purified by Qiaquick Gel Extraction Kit. The purified DNA was then used for standard library preparation and deep sequencing on an Illumina platform with 100 bp paired-end mode.

The MNase-seq data of *C. elegans* were analyzed as described by Li *et al*. (*28*). Briefly, the raw MNase-seq reads were mapped to the *C. elegans* genome (ce11) using BWA (version 0.7.17-r1188) (*41*), in which only uniquely-mapped reads were used for the following analyses. The mapped data were then subjected to DANPOS2 analysis (version 2.2.2) (*42*) for genome-wide profiling of nucleosome occupancy, with the strength of the MNase-seq signals calculated by deepTools (version 3.5.1) (*32*). The genomic regions of *C. elegans* were then divided into twenty bins across the genome, including ten bins of nucleosome-located regions and another ten bins of linker regions. The number of 6mA sites in each bin was calculated using BEDTools (v2.26.0) (*43*), and the 6mA level of each region was defined as the ratio of the number of the 6mA sites to the total number of A or T sites in this region.

### Histone modification analyses

*C. elegans* samples were treated with *P. aeruginosa* infection and collected as described above. The samples collected were suspended in ChIP cross-linking buffer (PBS with 1% formaldehyde) and rotated for 20 min. 2.5 M glycine (2/5 volume) was then added to stop the reaction. Worm pellet was washed using PBS buffer for three times, which was then resuspended in ChIP SDS lysis buffer (1% SDS, 10 mM EDTA, 50 mM Tris pH8.1) with PIC. The samples were then homogenized and incubated on ice for 20 min, following by sonicated with SONICS Vibra Cell sonicator. The supernatant was diluted with ChIP dilution buffer (0.01% SDS, 1.1% Triton X-100, 1.2 mM EDTA, 16.7 mM Tris-HCl pH8.1, 167 mM NaCl) after centrifugation, then 100 μL of pre-blocked Protein A agarose was added, following by rotation at 4°C for one hour. Then antibody was added to the supernatant separately (For H3K4me2: 4 μg per 30 μg chromatin; For H3K4me3: 4 μg per 30 μg chromatin; For H3K27me3: 6 μg per 30 μg chromatin), and rotated at 4°C for 16 hours. After the rotation, 25 μL of pre-blocked Dynabeads Protein G was added, and rotated at 4°C for 2 hours. The beads were washed with low-salt wash buffer (0.1% SDS, 1% Triton X- 100, 2 mM EDTA, 20 mM Tris-HCl pH8.1, 150 mM NaCl), high-salt wash buffer (0.1% SDS, 1% Triton X-100, 2 mM EDTA, 20 mM Tris-HCl pH8.1, 500 mM NaCl), and LiCl wash buffer (250 mM LiCl, 1% IGEPAL-CA630, 1% DOC). The beads were then wash with 1X TE again, and eluted with 250 μL elution buffer (1% SDS, 100 mM NaHCO3) for 20 min for two times. Finally, 20 μL of 5 M NaCl was added to the eluent and incubated at 65°C for 5 hours. 10 μL of 0.5 M EDTA, 20 μL of 1 M Tris-HCl pH6.5 and 20 μg proteinase K were then added, followed by the incubation at 45°C for two hours. The DNA obtained was purified by Zymo ChIP DNA Clean and Concentrator, which was further quantified by Qubit (Invitrogen) and examined by an Agilent 4200 for integrity. Finally, the wild-type *C. elegans* and METL-9 KO- OP50 *C. elegans* treated with H3K27me3, H3K4me2, or H3K4me3 antibody were then subjected to library preparation and deep sequencing on a MGI platform with 150 bp paired-end mode (Table S3).

For the ChIP-seq data, the raw sequencing reads were mapped to the *C. elegans* genome (ce11) using BWA (v0.7.13-r1126) with default parameters. Only properly paired, uniquely-mapped reads with mapping quality ≥ 20 and the number of mismatches < 8 were retained. PCR duplicates were removed by Picard (ver. 2.17.6), and the peak regions were obtained using Epic2 (version 0.0.52) (*44*). The strength of the ChIP-seq signals was calculated using deeptools (v2.4.2), by normalizing the reads coverage and subtracting the reads coverage at the corresponding region in the input sample.

## Acknowledgements

We thank Dr. Chuanhui Han and Dr. Shaokun Shu at Peking University for insightful suggestions. This work was supported by grants from the Ministry of Science and Technology of China (National Key Research and Development Program of China, 2018YFA0801405 and 2019YFA0801801), the National Natural Science Foundation of China (31871272) and the Chinese Institute for Brain Research (Beijing) (2020- NKX-XM-11).

## Author contributions

C.Y.L. and Y.L. conceived and designed the study. J.Z. and J.W. developed the new 6mA-Sniper method. Q.P. and C.M. performed most of the experiments. X.L., L.K.Z. performed part of the experiments. J.Z., Q.P., J,W., C.X., T.L., W.Z.Z., N.A.A., W.D. and L.Z. analyzed the data and performed statistical analysis. C.Y.L., J.Z. and Q.P. wrote the paper. All authors read and approved the final manuscript.

## Data and code availability

High-throughput sequencing data from this study have been submitted to the NCBI Sequence Read Archive (SRA) (https://www.ncbi.nlm.nih.gov/sra/) under accession number PRJNA857919. The codes in this study can be found at GitHub via URL: https://github.com/ZhangJiePKU/6mAProject.

## Supplementary information

This file contains Figure S1-S6 and Table S1-S3.

## Supplementary information

**Figure S1.**
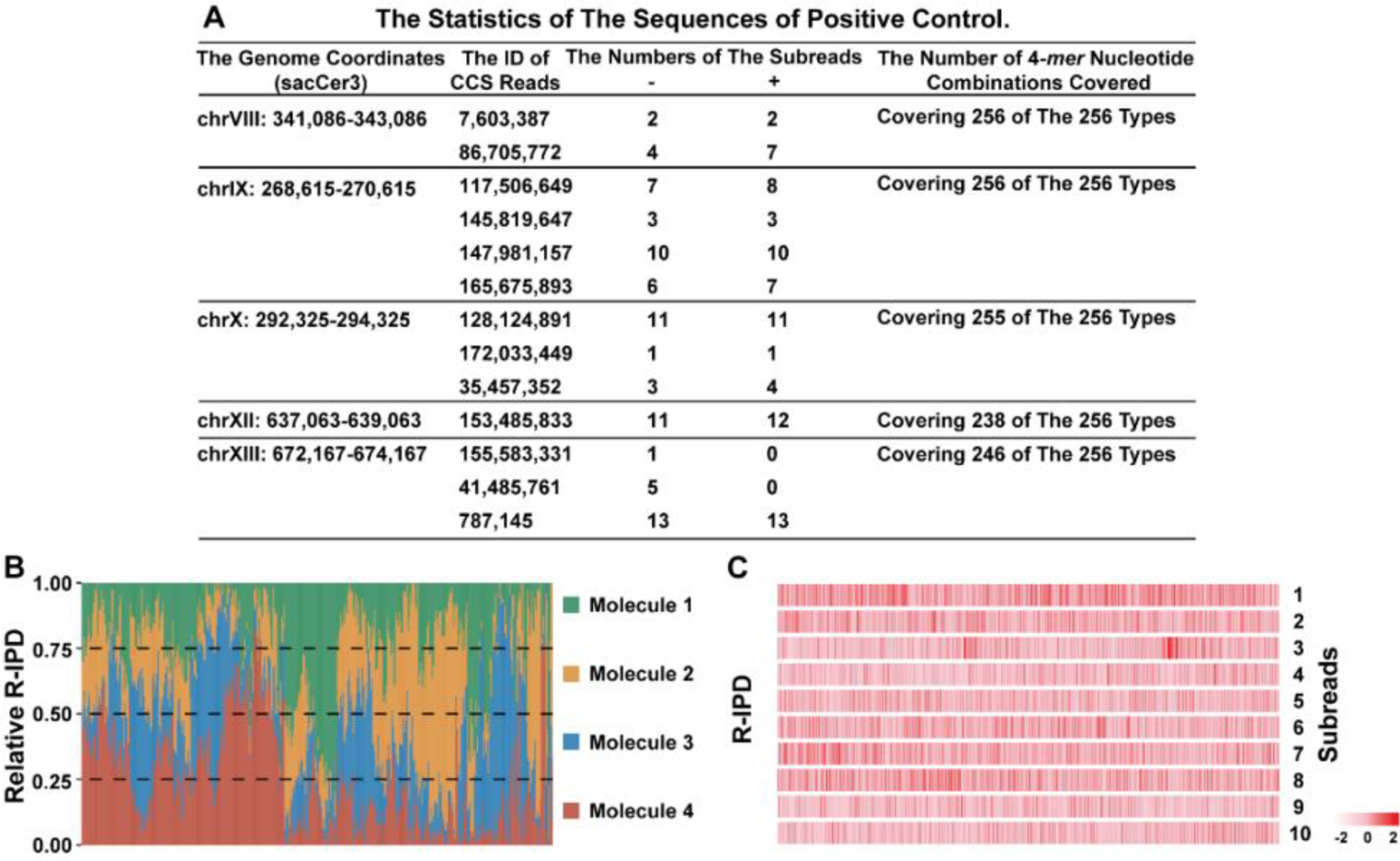
Statistics and features of the CCS reads. (**A**) Statistics for the sequences of positive control. Five yeast DNA sequences covering most of the 256 types of 4- *mer* nucleotide combinations were subjected to an *in vitro* methyltransferase assay, and the products were used as the positive control in 6mA identification. The genome coordinates (sacCer3), the ID of CCS reads, the numbers of the subreads on the plus and minus strand, and the number of 4-*mer* nucleotide combinations included were shown. (**B**) For these sequences, we selected the CCS reads of the PCR-amplified sample as the negative control. for each A site covered by four independent DNA molecules (chrIX: 268,617-270,510, one representative subreads of each molecule was randomly-selected), the raw IPD scores were normalized and shown. (**C**) For each site on one CCS reads (ID: 147,981,157), the raw IPD scores were normalized and shown in a heatmap according to the color scales, with the missing data shown in gray.

**Figure S2.**
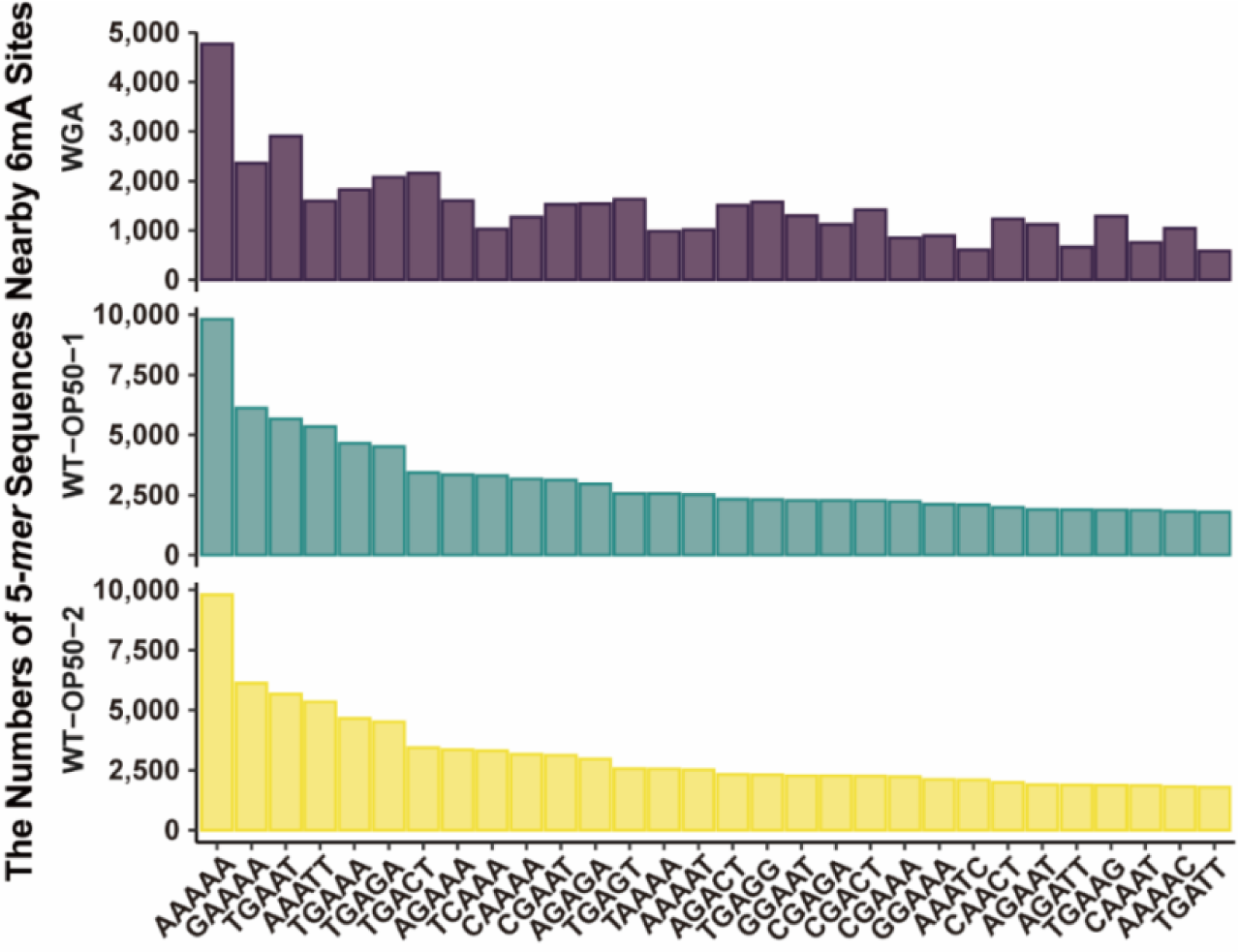
Motifs of candidate 6mA sites identified by the standard pipeline of SMRT sequencing. SMRT sequencing data from *C. elegans* (**WT-OP50-1**, **WT- OP50-2**, **WGA**) were analyzed using SMRT Link (v10.2) with default parameters to identify candidate 6mA sites. For each sample, the numbers of 5-*mer* sequences nearby these candidate 6mA sites were then counted and summarized.

**Figure S3.**
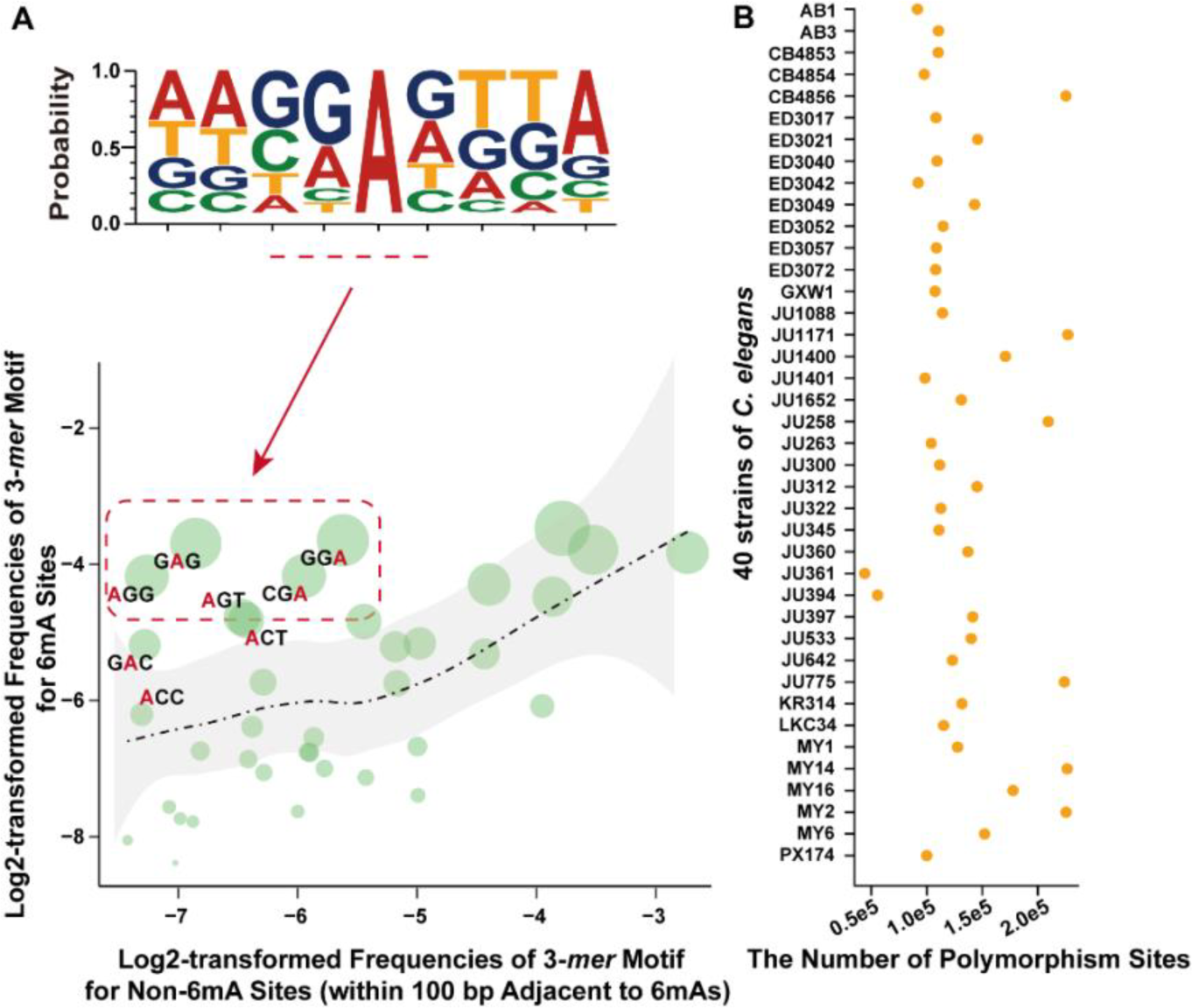
Identification of 6mA sites with a relaxed threshold. 10,692 6mA sites were identified in *C. elegans* genome, using a relaxed threshold (at least two, rather than three, positive CCS reads required in both of the samples). (**A**) The sequence motifs of these 6mA sites are shown, with the most significantly-enriched 3-*mer* sequences centered by 6mA sites highlighted in the red box. Each green bubble presents one of the various types of 3-*mer* nucleotide combinations with A sites. The size of each bubble represents the proportion of each 3-*mer* nucleotide combination, with the 6mA sites highlighted in red. The distribution of 3-*mer* nucleotide combination was estimated with random sequences, and shown in black dot line with gray regions representing the 95% confidence intervals. (**B**) In a population of 40 *C. elegans* strains, a total of 803,948 polymorphism sites were identified. The number of polymorphism sites for each strain was shown.

**Figure S4.**
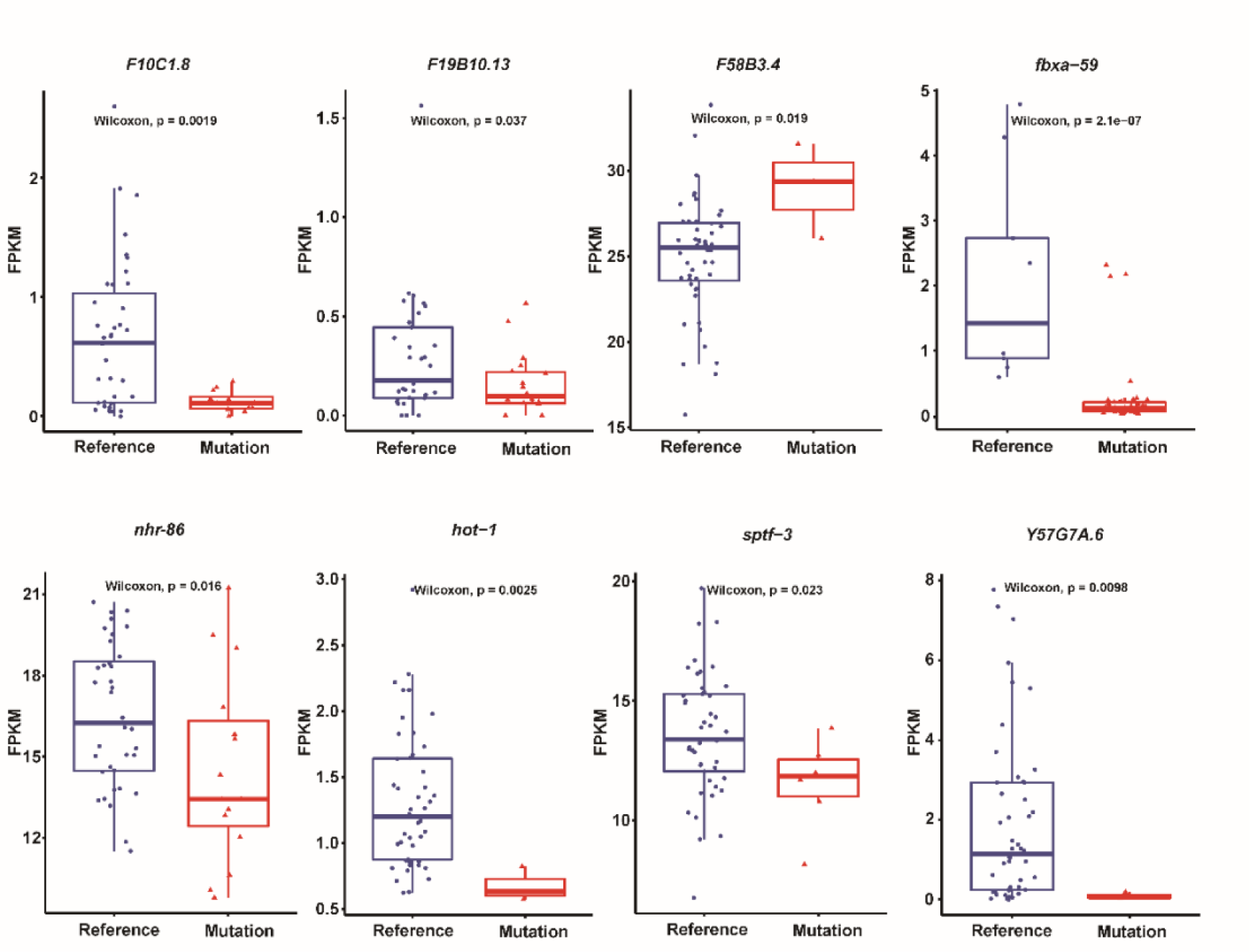
The distribution of the gene expression in *C. elegans* strains. For each polymorphic site co-localized with 6mA sites in the genic regions of the host gene, the expressions of the host gene were shown, for *C. elegans* strains with A allele at the polymorphic site (**Reference**), and the strains with other non-A alleles (**Mutation**). One-sided Wilcoxon tests were performed to test whether the expression levels between the two groups are significantly different.

**Figure S5.**
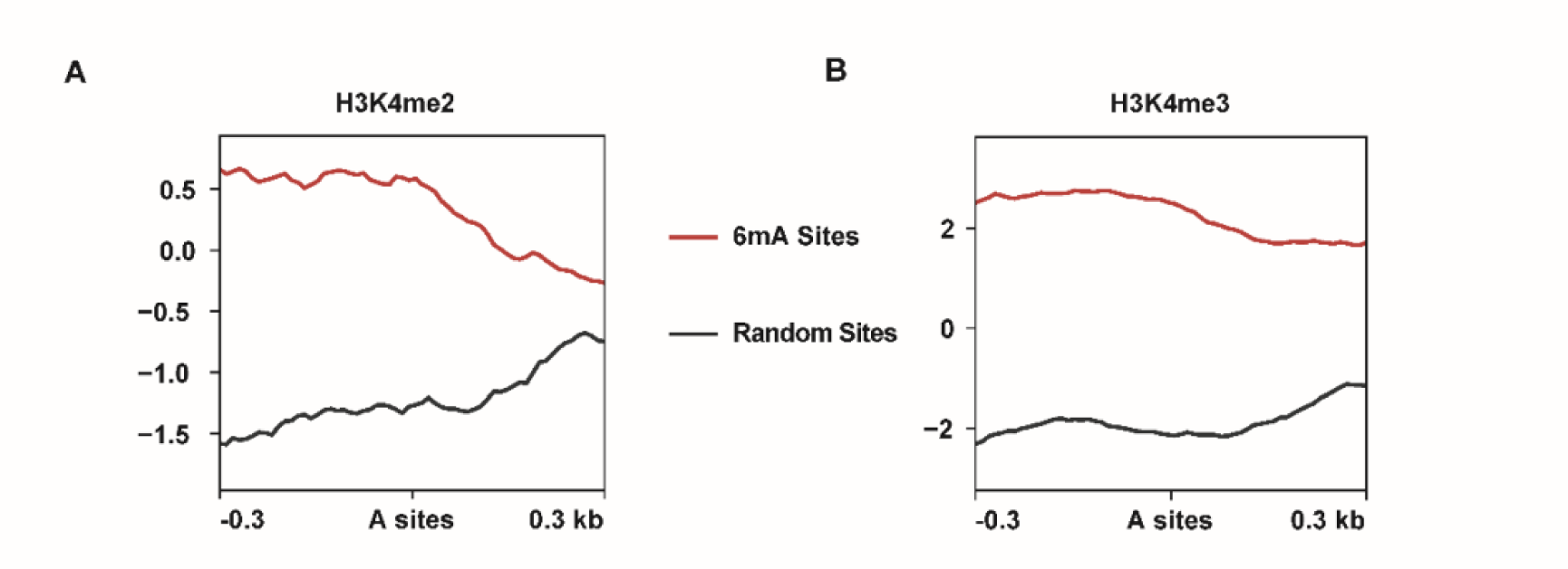
H3K4me2 and H3K4me3 signals at 6mA-marked regions. (**A-B**) For the focal 6mA sites and their surrounding regions, the levels of H3K4me2 (**A**) and H3K4me3 (**B**) were shown in red lines. As a background, the levels of H3K4me2 (**A**) and H3K4me3 (**B**) were also shown in black lines, for randomly-selected A sites and their surrounding regions.

**Figure S6.**
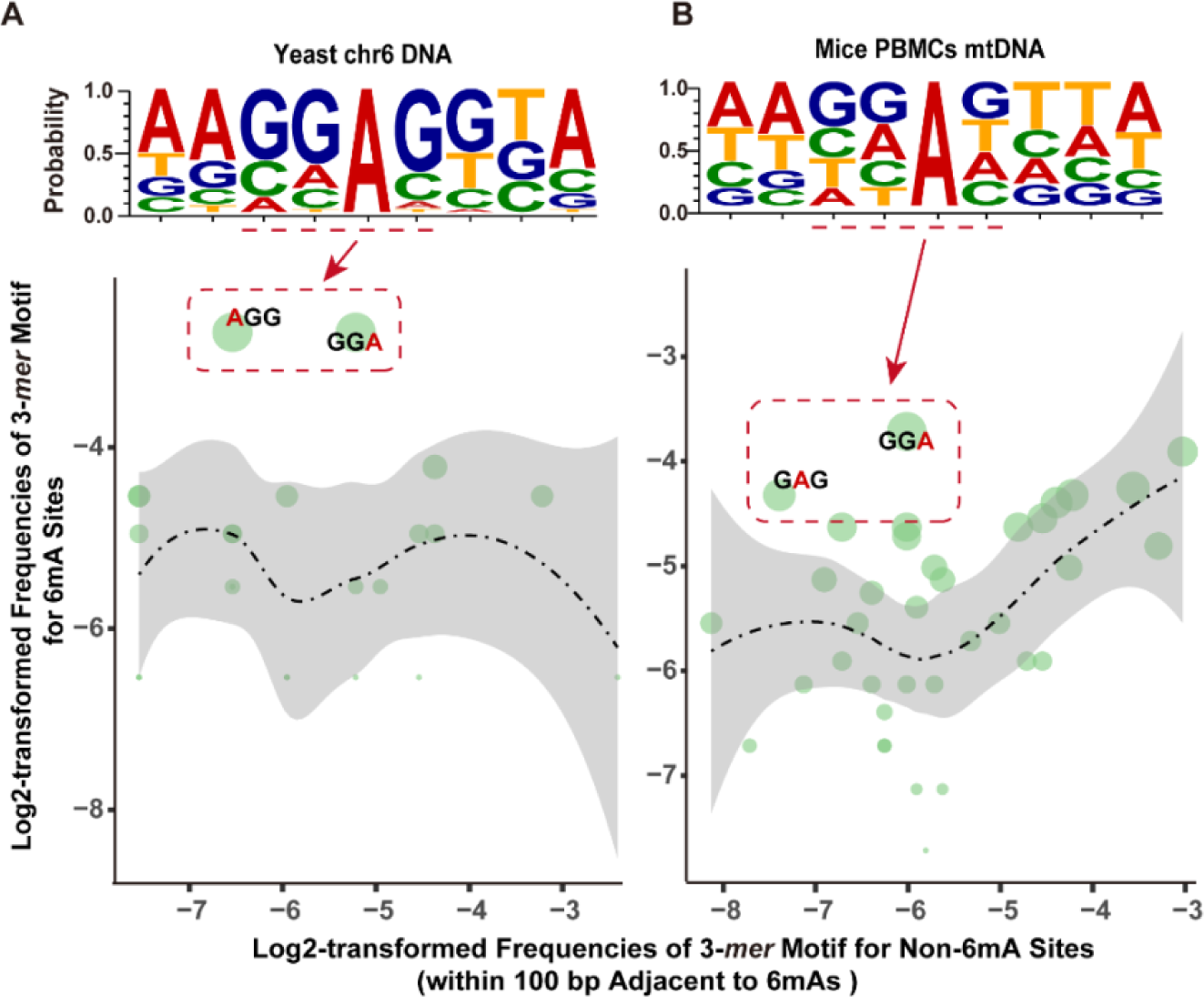
Identification of 6mA events in yeast and mouse. (**A-B**) 31 6mA events on chromosome 6 in yeast, and 142 events on mitochondria genomes (mtDNA) in PBMCs of mice were identified. The sequence motifs of these 6mA sites in yeast (**A**) and mouse (**B**) are shown, with the most enriched 3-*mer* sequences centered by 6mA sites highlighted in red boxes. Each green bubble presents one of the various types of 3-*mer* nucleotide combinations with A sites. The size of each bubble represents the proportion of each 3-*mer* nucleotide combination, with the 6mA sites highlighted in red. The distribution of 3-*mer* nucleotide combination was estimated with random sequences, and shown in black dot line with gray regions representing 95% confidence intervals.

**Table S1.** | Statistics of the whole-genome sequencing data of *C. elegans*, yeast and mouse.

| Species | Samples | Total Reads | Mapped Reads | Total Mapping Rate (%) |
| --- | --- | --- | --- | --- |
| *C. elegans* | WT-OP50-1 | 9,576,822 | 9,468,525 | 98.8 |
|  | WT-OP50-2 | 12,615,770 | 12,468,466 | 98.8 |
|  | WT-PA14-1 | 4,489,023 | 4,446,396 | 99.1 |
|  | WT-PA14-2 | 5,243,813 | 5,198,392 | 99.1 |
|  | KO-OP50-1 | 13,185,298 | 13,028,127 | 98.8 |
|  | KO-OP50-2 | 12,366,594 | 12,242,837 | 98.9 |
|  | KO-PA14-1 | 2,762,815 | 2,722,964 | 98.5 |
|  | KO-PA14-2 | 3,008,663 | 2,968,038 | 98.6 |
|  | WGA-1 | 7,818,207 | 6,556,326 | 83.8 |
| Yeast | WT-1 | 25,253,602 | 24,650,874 | 97.6 |
|  | WGA-1 | 15,721,081 | 11,484,072 | 73.0 |
| Mouse | WT-1 | 187,749,588 | 182,312,327 | 97.1 |
|  | WGA-1 | 95,388,422 | 897,270,69 | 94.1 |

**Table S2.**
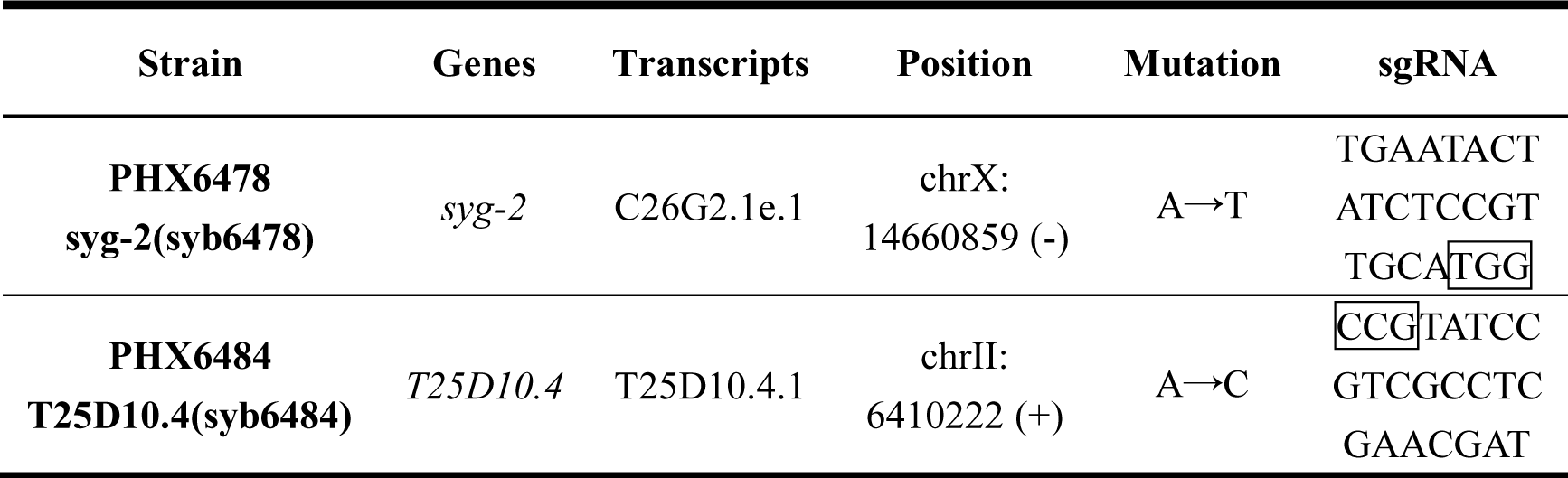
| Information of mutated *C. elegans* strains.

**Table S3.** | Statistics of the ChIP-seq data of H3K27me3/H3K4me2/H3K4me3.

| **Histone**<br>**modification** | **Strain** | **Total**<br>**Reads** | **Mapped**<br>**Reads** | **Total**<br>**Mapping**<br>**Rate (%)** | **Unique**<br>**Mapped**<br>**Reads** | **Unique**<br>**Mapping**<br>**Rate (%)** |
| --- | --- | --- | --- | --- | --- | --- |
| **H3K27me3** | WT-OP50 | 12,699,413 | 8,633,386 | 67.98 | 7,561,648 | 59.5 |
|  | METL-9<br>KO-OP50 | 11,032,171 | 9,698,241 | 87.91 | 8,537,315 | 77.4 |
| **H3K4me2** | WT-OP50 | 23,546,583 | 23,505,303 | 99.82 | 21,801,061 | 92.6 |
|  | METL-9<br>KO-OP50 | 46,427,836 | 46,299,769 | 99.72 | 43,379,514 | 93.4 |
| **H3K4me3** | WT-OP50 | 23,194,565 | 22,462,363 | 96.84 | 20,502,931 | 88.4 |
|  | METL-9<br>KO-OP50 | 23,236,511 | 21,973,851 | 94.57 | 20,090,986 | 86.5 |

## References

1. Z. D. Smith, A. Meissner, DNA methylation: roles in mammalian development. Nature Reviews Genetics 14, 204–220 (2013).

2. J. A. Law, S. E. Jacobsen, Establishing, maintaining and modifying DNA methylation patterns in plants and animals. Nature Reviews Genetics 11, 204–220 (2010).

3. Z. Hao et al., N6-Deoxyadenosine Methylation in Mammalian Mitochondrial DNA. Molecular Cell 78, 382–395.e388 (2020).

4. K. Douvlataniotis, M. Bensberg, A. Lentini, B. Gylemo, C. E. Nestor, No evidence for DNA N-6-methyladenine in mammals. Sci Adv 6 (2020).

5. D. B. Dunn, J. D. Smith, Occurrence of a New Base in the Deoxyribonucleic Acid of a Strain of Bacterium-Coli. Nature 175, 336–337 (1955).

6. Z. Li et al., N6-methyladenine in DNA antagonizes SATB1 in early development. Nature 583, 625–630 (2020).

7. Y. Fu et al., N6-Methyldeoxyadenosine Marks Active Transcription Start Sites in Chlamydomonas. Cell 161, 879–892 (2015).

8. L. Y. Beh et al., Identification of a DNA N6-Adenine Methyltransferase Complex and Its Impact on Chromatin Organization. Cell 177, 1781–1796.e1725 (2019).

9. G. Zhang et al., N6-Methyladenine DNA Modification in Drosophila. Cell 161, 893–906 (2015).

10. T. P. Wu et al., DNA methylation on N6-adenine in mammalian embryonic stem cells. Nature 532, 329–333 (2016).

11. K. Boulias, E. L. Greer, Means, mechanisms and consequences of adenine methylation in DNA. Nature Reviews Genetics 23, 411–428 (2022).

12. Z. K. O’Brown et al., Sources of artifact in measurements of 6mA and 4mC abundance in eukaryotic genomic DNA. BMC Genomics 20 (2019).

13. Y. M. Kong et al., Critical assessment of DNA adenine methylation in eukaryotes using quantitative deconvolution. Science 375, 515 (2022).

14. G.-Z. Luo et al., Characterization of eukaryotic DNA N6-methyladenine by a highly sensitive restriction enzyme-assisted sequencing. Nature Communications 7 (2016).

15. A. M. Wenger et al., Accurate circular consensus long-read sequencing improves variant detection and assembly of a human genome. Nature Biotechnology 37, 1155–1162 (2019).

16. J. J. Kasianowicz, S. M. Bezrukov, On ’three decades of nanopore sequencing’. Nature Biotechnology 34, 481–482 (2016).

17. Eric L. Greer et al., DNA Methylation on N6-Adenine in C. elegans. Cell 161, 868–878 (2015).

18. C. Ma et al., N6-methyldeoxyadenine is a transgenerational epigenetic signal for mitochondrial stress adaptation. Nature Cell Biology 21, 319–327 (2018).

19. Q. L. Wan et al., N-6-methyldeoxyadenine and histone methylation mediate transgenerational survival advantages induced by hormetic heat stress. Sci Adv 7 (2021).

20. X. Liu et al., N6-methyladenine is incorporated into mammalian genome by DNA polymerase. Cell Research 31, 94–97 (2020).

21. X. Li et al., The exploration of N6-deoxyadenosine methylation in mammalian genomes. Protein & Cell 12, 756–768 (2021).

22. M. U. Musheev, A. Baumgärtner, L. Krebs, C. Niehrs, The origin of genomic N6-methyl- deoxyadenosine in mammalian cells. Nature Chemical Biology 16, 630–634 (2020).

23. P. Spingardi, S. Kriaucionis, How m6A sneaks into DNA. Nature Chemical Biology 16, 604–605 (2020).

24. L.-Q. Chen et al., High-precision mapping reveals rare N6-deoxyadenosine methylation in the mammalian genome. Cell Discovery 8 (2022).

25. C. Ma, T. Xue, Q. Peng, J. Zhang, J. Guan, W. Ding, Y. Li, P. Xia, L. Zhou, T. Zhao, S. Wang, L. Quan, C. Y. Li, Y. Liu. N6-Deoxyadenine Methyltransferase Modulates C. elegans Immunity via Dichotomous Mechanisms. Cell Research, revision (2023).

26. Y. Duan et al., Linkage of A-to-I RNA Editing in Metazoans and the Impact on Genome Evolution. Molecular Biology and Evolution 35, 132–148 (2018).

27. R. K. Chodavarapu et al., Relationship between nucleosome positioning and DNA methylation. Nature 466, 388–392 (2010).

28. Y. Li et al., Human exonization through differential nucleosome occupancy. Proceedings of the National Academy of Sciences 115, 8817–8822 (2018).

29. Y. Li et al., Replication-Independent Histone Turnover Underlines the Epigenetic Homeostasis in Adult Heart. Circulation Research 125, 198–208 (2019).

30. G. E. Crooks, G. Hon, J. M. Chandonia, S. E. Brenner, WebLogo: A sequence logo generator. Genome Research 14, 1188–1190 (2004).

31. J. Jänes et al., Chromatin accessibility dynamics across C. elegans development and ageing. eLife 7 (2018).

32. F. Ramírez, F. Dündar, S. Diehl, B. A. Grüning, T. Manke, deepTools: a flexible platform for exploring deep-sequencing data. Nucleic Acids Research 42, W187–W191 (2014).

33. O. Thompson et al., The million mutation project: A new approach to genetics in Caenorhabditis elegans. Genome Research 23, 1749–1762 (2013).

34. D. Kim, J. M. Paggi, C. Park, C. Bennett, S. L. Salzberg, Graph-based genome alignment and genotyping with HISAT2 and HISAT-genotype. Nature Biotechnology 37, 907–915 (2019).

35. H. Li et al., The Sequence Alignment/Map format and SAMtools. Bioinformatics 25, 2078–2079 (2009).

36. A. Tarasov, A. J. Vilella, E. Cuppen, I. J. Nijman, P. Prins, Sambamba: fast processing of NGS alignment formats. Bioinformatics 31, 2032–2034 (2015).

37. Y. Liao, G. K. Smyth, W. Shi, featureCounts: an efficient general purpose program for assigning sequence reads to genomic features. Bioinformatics 30, 923–930 (2014).

38. M. D. Robinson, D. J. McCarthy, G. K. Smyth, edgeR: a Bioconductor package for differential expression analysis of digital gene expression data. Bioinformatics 26, 139–140 (2010).

39. M. I. Love, W. Huber, S. Anders, Moderated estimation of fold change and dispersion for RNA- seq data with DESeq2. Genome Biology 15 (2014).

40. G. Zhang, N. M. Roberto, D. Lee, S. R. Hahnel, E. C. Andersen, The impact of species-wide gene expression variation on Caenorhabditis elegans complex traits. Nature Communications 13 (2022).

41. H. Li, R. Durbin, Fast and accurate short read alignment with Burrows–Wheeler transform. Bioinformatics 25, 1754–1760 (2009).

42. K. Chen et al., DANPOS: Dynamic analysis of nucleosome position and occupancy by sequencing. Genome Research 23, 341–351 (2013).

43. A. R. Quinlan, I. M. Hall, BEDTools: a flexible suite of utilities for comparing genomic features. Bioinformatics 26, 841–842 (2010).

44. E. B. Stovner, P. Saetrom, epic2 efficiently finds diffuse domains in ChIP-seq data. Bioinformatics 35, 4392–4393 (2019).

